# Reinforcement sculpts synaptic ensembles for operant dendritic control

**DOI:** 10.64898/2026.11.04.764390

**Authors:** Eddy Albarran, Akira Fushiki, Vivek Athalye, Darcy S. Peterka, Rui M. Costa

**Affiliations:** Zuckerman Institute, Columbia University, New York, NY; Allen Institute, Seattle, WA

## Abstract

Reinforcement learning requires the selective modification of synapses that contribute to outcome-relevant neural activity, yet the principles underlying these selective modifications remain unclear. Here we combined longitudinal two-photon imaging of identified dendritic spines with a closed-loop optical brain–machine interface (BMI), in which activity from selected apical dendrites of layer 5 motor-cortical neurons controlled an auditory cursor, with target acquisition triggering optogenetic reinforcement. Mice learned to reliably generate the specific dendritic activity that leads to reinforcement. Learning was accompanied by selective strengthening and weakening of individual spines on reinforced dendrites. At individual synapses, the direction of plasticity depended on both the timing of activity relative to reinforced dendritic events and the pre-existing temporal organization of that activity. Furthermore, plasticity was enriched in small task-relevant synaptic groups: spines in synergistic groups were preferentially strengthened, whereas those in antagonistic groups were preferentially weakened. These changes were accompanied by the emergence of increasingly reproducible, selectively recruited, and temporally focused synaptic ensembles, and the incorporation of newly formed spines into these ensembles. Finally, blocking striatal dopamine receptor signaling impaired BMI learning and cortical synaptic reorganization of target dendrites, demonstrating a role for striatal reinforcement in cortical plasticity. Together, these findings reveal how reinforcement of specific dendritic activity sculpts synaptic ensembles that support learned operant dendritic control.

## Main text

Learning requires changes at particular synapses among the thousands of inputs received by individual cortical neurons. Which synapses are modified during reinforcement learning, and what governs the direction and magnitude of their changes, remain difficult to establish in vivo. Synapses active near a successful neural event need not all contribute equally to that event, raising a fundamental question of how reinforcement becomes associated with lasting, input-specific plasticity^1–4^.

In conventional motor learning tasks, movements and their consequences unfold alongside sensory feedback, ongoing neural activity, and interactions among distributed cortical and subcortical circuits^5–8^. Identifying the neural substrates of learning therefore presents a fundamental challenge: among the large number of neurons engaged during behavior, it is difficult to determine which are relevant to successful outcomes and which of their vast synaptic inputs are selected for reinforcement-dependent plasticity, akin to finding a needle in a haystack. Experimentally defining which cortical neurons lead to reinforcement offers a way to narrow this search and investigate synaptic selection within an identified neural population.

Optical brain-machine interfaces (BMIs) provide this experimental leverage by linking the activity of selected cortical populations to a feedback signal and a precisely defined reinforcement contingency^9–16^. Here we extended this paradigm to dendrites and synapses by developing a closed-loop optical BMI in which activity from specific apical dendrites of layer 5 motor-cortical neurons triggered reinforcement via optogenetic stimulation of dopamine neurons in the ventral tegmental area (VTA). Longitudinal imaging of identified dendritic spines in these dendrites allowed us to measure synaptic activity around the events that lead to reinforcement and follow changes in those same inputs over learning.

We found that reinforcement-associated synaptic strengthening and weakening depended not simply on coincident synaptic activity, but on the timing of recruitment relative to reinforced dendritic events and also the pre-existing temporal organization. Further, plasticity was preferentially observed in small synaptic ensembles encoding task-relevant information. These selective changes were concomitant with the emergence of increasingly reproducible and temporally focused synaptic ensembles. Finally, dopamine receptor blockade in the striatum impaired both BMI learning and synaptic reorganization in the cortex. Together, these findings demonstrate that animals can learn to operantly control the activity of specific cortical dendrites and reveal novel principles of synaptic plasticity that enable such learning and control.

## Results

### Learned control of specific cortical dendritic activity through an optical BMI

To investigate how reinforcement shapes cortical activity and synaptic plasticity, we developed a closed-loop optical BMI that coupled the activity of identified cortical dendrites to optogenetic reinforcement (Fig. 1a–c). We selectively imaged corticospinal pyramidal neurons in layer 5 of primary motor cortex (M1) of DAT-FlpO mice, using retrograde viral targeting of the cervical spinal cord, resulting in sparse expression of GCaMP8m and tdTomato to enable functional and structural imaging of apical dendrites and dendritic spines (Fig. 1b,h). Apical dendritic segments of two imaged neurons (E1 and E2) controlled an auditory cursor in opposing directions, with cursor movement to the specified E1 target (hit) triggering closed-loop optogenetic stimulation of ChR2-expressing dopamine neurons in the VTA (Methods). E2-directed activity was not reinforced. We repeatedly imaged the same identified dendrites and their associated dendritic spines across baseline and eight consecutive training sessions, allowing us to longitudinally track apical dendritic and spine Ca^2+^ activity throughout learning.

**Figure 1.**
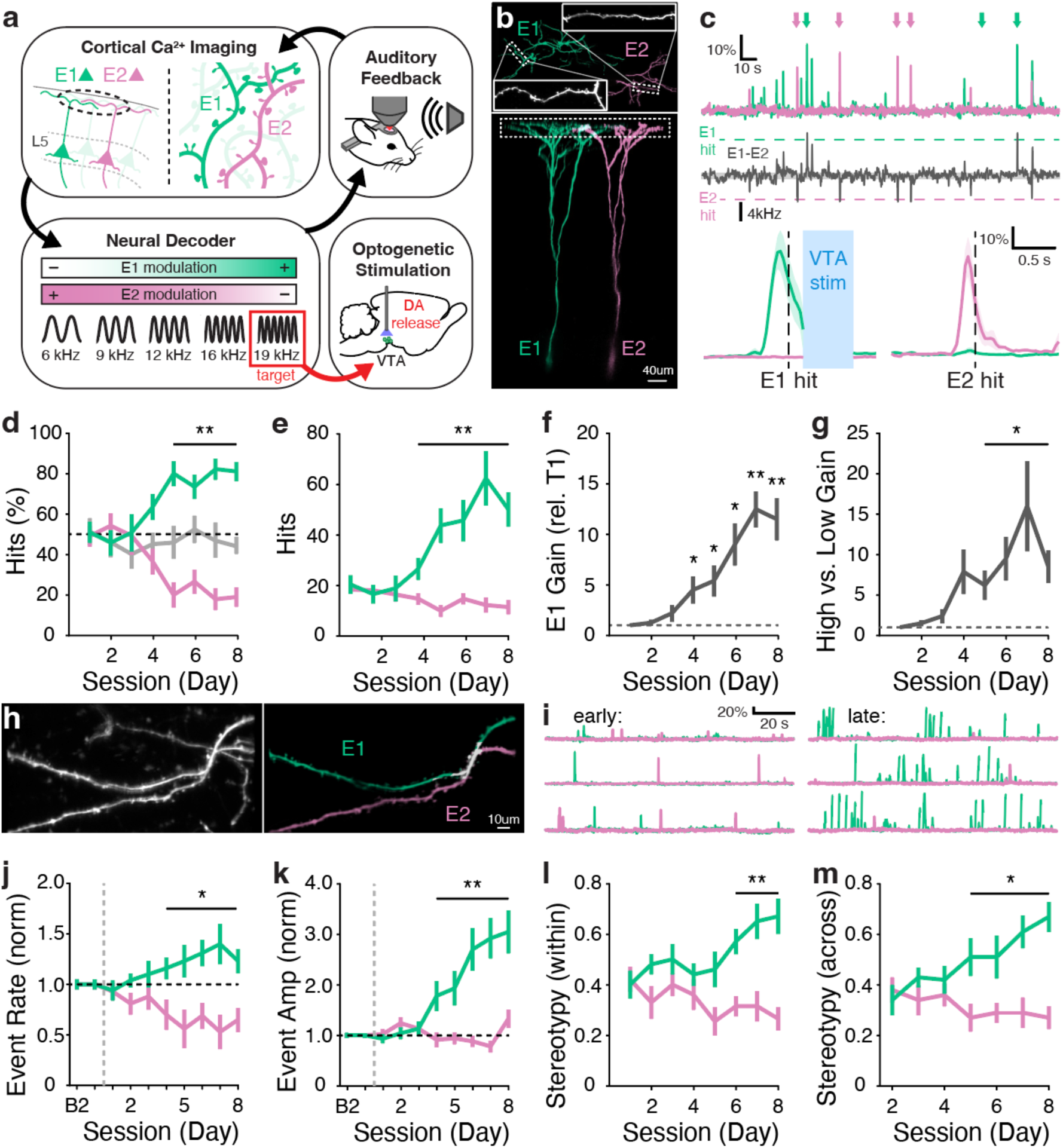
Learned control of specific cortical dendritic activity through an optical BMI. a, Schematic of the closed-loop optical brain-machine interface (BMI). Two populations of imaged cortical dendrites (E1 and E2) control an auditory cursor through their differential calcium activity. Cursor movement toward the E1 target triggers optogenetic stimulation of channelrhodopsin-2 (ChR2)-expressing ventral tegmental area (VTA) dopamine neurons, whereas E2 target acquisition is not reinforced. b, Representative two-photon images of sparsely labeled layer 5 corticospinal neurons and their apical dendrites, illustrating the imaged E1 (green) and E2 (magenta) populations. Insets show representative dendritic segments. Scale bar, 40 μm. c, Representative E1 and E2 dendritic calcium activity (top), decoded E1–E2 cursor signal and target thresholds (middle), and event-aligned calcium responses during E1 and E2 target acquisition (bottom). Optogenetic VTA stimulation (0.6 s) follows E1 target acquisition. d, Fraction of E1 and E2 target hits across training sessions, shown alongside YFP control animals (grey). e, Number of E1 and E2 target hits across training sessions. f, Gain in E1-directed cursor occupancy relative to the first training session (T1). g, Relative gain in high-versus low-E1 cursor occupancy across training. h, Representative two-photon images of identified E1 and E2 dendritic segments. Scale bar, 10 μm. i, Representative E1 and E2 dendritic calcium activity during early and late training (T2 and T7 in this example), illustrating the increasing prevalence of E1-associated events. j,k, Baseline-normalized dendritic calcium-event rate (j) and amplitude (k) across baseline and training sessions. l,m, Within-session (l) and across-session (m) stereotypy of E1 and E2 cursor trajectories during training. Green, E1; magenta, E2; grey, YFP controls where indicated. n = 6 ChR2-VTA mice, 5 YFP-VTA mice. Data are presented as mean ± s.e.m..

Mice progressively acquired selective control of the reinforced E1 target (Fig. 1d,e). Following an initial period of variable target-directed performance, E1 hit rates increased markedly during training and remained elevated in subsequent sessions. Consistent with this improvement, cursor trajectories exhibited increasing occupancy toward the reinforced E1 target (Fig. 1f,g). This performance progression defined three learning epochs: early training (T1–T3), a transition phase (T4–T5) characterized by rapid acquisition of E1 control, and late training (T6–T8), during which performance remained elevated. Reinforced E1 cursor trajectories also became increasingly stereotyped within and across sessions, whereas E2 trajectories showed no comparable stabilization (Fig. 1l,m; Extended Data Fig. 1a), consistent with the emergence of a reproducible cortical solution for reaching the reinforced target. Analyses using a common baseline-defined cursor calibration confirmed that these changes were not explained by session-specific normalization (Extended Data Fig. 1b,c).

Notably, BMI learning was accompanied by a pronounced reorganization of E1 and E2 dendritic activity (Fig. 1i), with selective increases in E1 dendritic event rates and amplitudes during training (Fig. 1j,k). Furthermore, E1 dendritic events became increasingly clustered in time as learning progressed, while direct coupling between E1 and E2 dendritic activity decreased (Extended Data Fig. 1d-g). A separate cohort of mice expressing YFP in VTA DA neurons did not exhibit increases in E1 hit rates or changes in dendritic activity dynamics, suggesting that the observed changes in ChR2-expressing VTA mice are driven by a dopamine signaling mechanism (Fig. 1d; Extended Data Fig. 1h-j). Together, these results indicate that BMI reinforcement establishes increasingly reliable generation of reinforced dendritic activity patterns.

### Synaptic activity exhibits structured temporal dynamics during baseline and reinforcement

To investigate the activity dynamics of individual synapses during reinforcement learning, we longitudinally tracked identified dendritic spines across baseline and training sessions (Fig. 2a–d). We isolated spine-local calcium transients from their parent dendritic signals (Methods) and quantified their event rates, amplitudes, and timing relative to dendritic activity. All analyses in this study focused on spine events that were well isolated from dendritic contamination (see Methods)^17–20^. Local infusion of the NMDA receptor antagonist R-CPP markedly suppressed these isolated events, consistent with a predominant origin of NMDAR-dependent synaptic activity (Extended Data Fig. 2a). Despite the pronounced changes in E1 dendritic activity accompanying BMI learning, population-average spine event rates and amplitudes remained largely unchanged across training (Fig. 2e,f). Thus, BMI learning was not accompanied by a gross increase in synaptic activity across the reinforced E1 dendrite.

**Figure 2.**
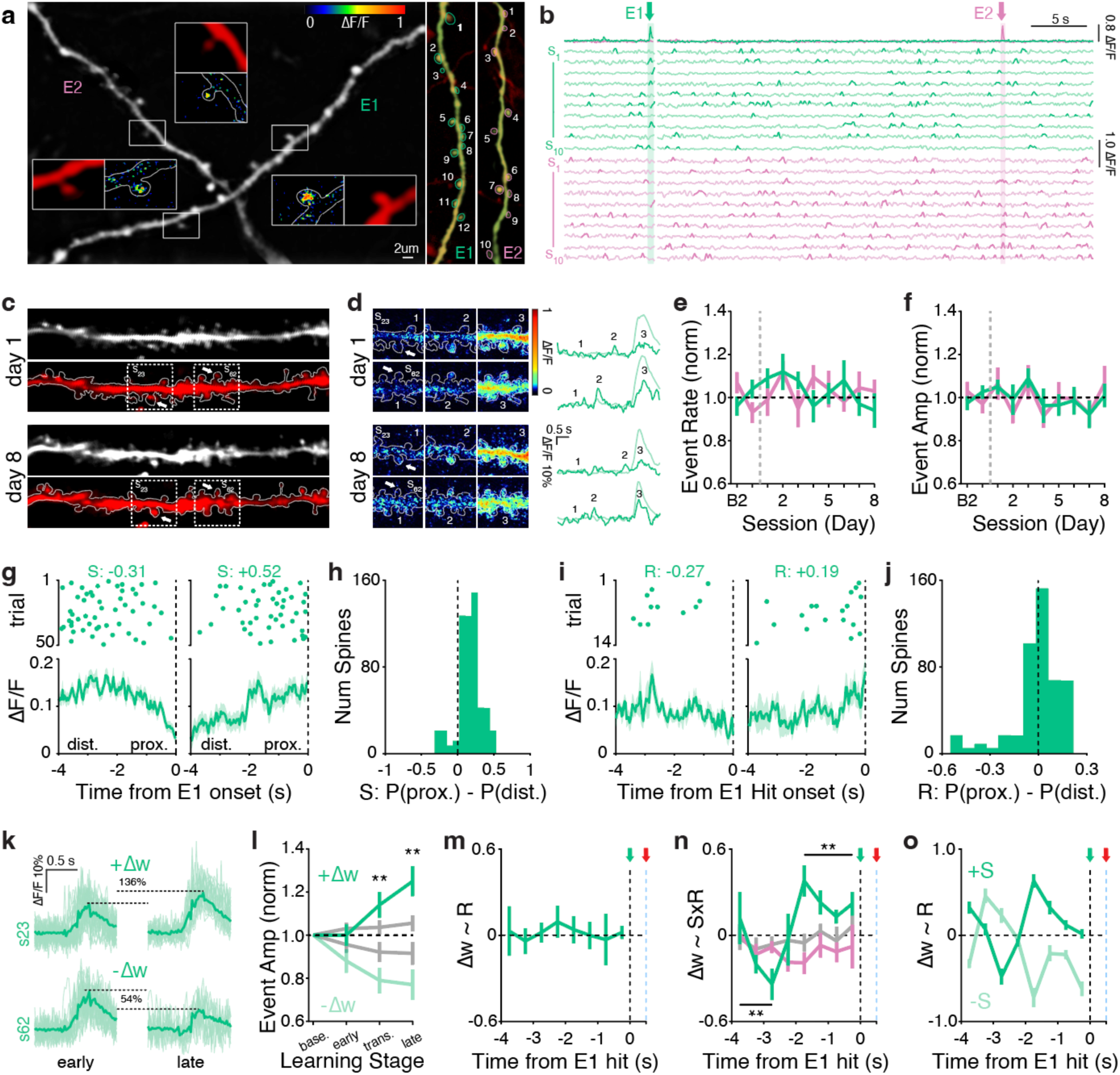
Reinforcement gates plasticity according to pre-existing synaptic organization and recruitment timing. a, Representative imaging session illustrating identification of dendritic spines on E1 and E2 dendrites. Insets show structural fluorescence, spine region-of-interest segmentation and examples of isolated spine calcium signals. Representative E1 and E2 dendritic segments and identified spines are shown at right. Scale bar, 2 μm. b, Simultaneous calcium activity from identified E1 (green) and E2 (magenta) dendritic spines, aligned with representative E1 and E2 target hits (arrows). c, Representative longitudinal imaging of the same dendritic segment during early (day 1) and late (day 8) training, illustrating identification and tracking of individual spines (arrows: left, s23; right, s62). d, Example spine-local calcium events measured in identified spines across training sessions, including spatial maps of calcium activity (dotted regions in (c)) and corresponding fluorescence traces. e,f, Baseline-normalized isolated spine calcium-event rate (e) and amplitude (f) across baseline and training for E1 and E2 spine populations. g, Representative baseline event raster plots and dendrite-aligned activity profiles for two spines (left-right) illustrating distinct temporal recruitment patterns preceding E1 dendritic-event onset (dotted vertical line). *S* describes the relative probability of temporally proximal versus distal recruitment. h, Distribution of baseline *S* values across identified E1 spines. i, Representative event raster plots and activity profiles for two spines (left-right) illustrating differences in recruitment preceding E1 hits. *R* describes relative recruitment during proximal versus distal portions of the pre-event interval. j, Distribution of *R* values across E1 spines. k, Representative longitudinal spine calcium responses illustrating two spines that increased (s23; +Δw, 136%) or decreased (s62; −Δw, 54%) event amplitudes during learning. l, Changes in normalized isolated spine-event amplitude across learning stages for strengthened (+Δw) and weakened (−Δw) spines, with YFP controls shown in grey. m, Time-resolved relationship between subsequent plasticity (Δw) and reinforcement-associated spine recruitment (*R*) preceding successful E1 dendritic events. n, Time-resolved relationship between Δw and the interaction between baseline temporal state and reinforcement-associated recruitment (*S* × *R*), shown for ChR2 and YFP conditions. o, Relationship between Δw and reinforcement-associated recruitment (*R*) stratified by positive and negative baseline temporal state (*S*). The vertical dashed line denotes dendritic-event onset; the additional arrow markers denote E1 hit detection (green) and optical stimulation (red). Green, E1 or strengthened spines as indicated; pale green, weakened spines; magenta, E2; grey, YFP controls. n = 768 spines / 12 dendrites / 6 mice (64 ± 3.24 spines / dendrite). Data are presented as mean ± s.e.m..

We next examined the temporal dynamics of synaptic activity preceding E1 dendritic events. During baseline sessions, spine activity exhibited a reproducible temporal structure spanning several seconds before dendritic-event onset (Fig. 2g,h). Individual spines differed in their probability of participating at different times within this dendrite-centered sequence. We summarized this pre-existing temporal organization using a spine-specific temporal-state coordinate, *S*, reflecting the relative probability of recruitment during temporally proximal versus distal portions of the sequence.

We then examined how individual spines were recruited during reinforced E1 dendritic events. Spine event times remained temporally heterogeneous, with different spines exhibiting distinct distributions of activity preceding reinforcement-associated dendritic events (Fig. 2i,j). We quantified these differences using a reinforcement-timing variable, *R*, describing each spine’s relative recruitment during proximal versus distal periods preceding reinforced dendritic events. Thus, individual synapses exhibited distinct temporal organizations both before learning and during reinforcement.

### Reinforcement gates plasticity according to pre-existing synaptic organization and recruitment timing

Longitudinal measurements revealed substantial heterogeneity in synaptic plasticity, with individual spines undergoing persistent strengthening (+Δw) or weakening (−Δw) in median event sizes relative to baseline (Fig. 2k,l). To identify potential determinants of these divergent outcomes, we first examined several candidate predictors. Neither baseline spine event amplitudes nor event rates reliably predicted subsequent plasticity (Extended Data Fig. 2b). Surprisingly, plasticity was also not predicted by the frequency with which individual spines participated in reinforced dendritic events (Fig. 2m). These findings argue against simple models in which the most active, strongest, or even most frequently reinforced cortical synapses are preferentially strengthened during reinforcement learning.

Instead, synaptic plasticity was strongly predicted by the interaction between each spine’s pre-existing temporal organization (*S*) and its temporal recruitment during reinforced dendritic events (*R*) (Fig. 2n,o). This relationship was most pronounced during the transition stage of learning (T4-T5), coinciding with the emergence of reliable E1 target acquisition, and was comparatively weak during early and late training (Extended Data Fig. 2c). Time-resolved analyses localized the plasticity rule to the seconds preceding reinforced dendritic-event onset: reinforcement-associated activity within an approximately 1.6-s window preferentially strengthened those inputs with a prior (baseline) bias toward dendrite-proximal recruitment (positive *S*), whereas activity occurring more than approximately 2-s before reinforced E1 events was associated with subsequent weakening (Fig. 2n,o).

Importantly, control animals expressing YFP rather than ChR2 in VTA dopamine neurons did not exhibit bidirectional plasticity or the corresponding temporal plasticity relationship observed during optogenetic reinforcement (Fig. 2l,n; grey). Furthermore, local cortical infusion of R-CPP during training blocked the divergence of +Δw and −Δw spine populations and impaired BMI learning (Extended Data Fig. 2d,e). In a separate cohort, we additionally imaged synaptic glutamate signals using iGluSnFR3 combined with jRGECO, but unlike spine calcium measurements, glutamate-defined Δw did not exhibit significant reinforcement-dependent divergence relative to YFP controls (Extended Data Fig. 2f-h).

Together, these findings identify a reinforcement-gated temporal plasticity rule in which the consequences of synaptic recruitment depend on the pre-existing temporal organization of individual inputs. The same reinforcement signal can therefore be associated with strengthening or weakening at different synapses, depending on their position within the pre-existing dendrite-centered temporal landscape.

### Plasticity is enriched in synergistic task-relevant synaptic groups

In addition to their individual temporal organization, synaptic inputs can be spatiotemporally integrated within local dendritic compartments^21–24^, raising the possibility that their functional interactions are also related to plasticity. We first examined the spatial organization of E1 spine activity. Nearby spines, particularly those separated by less than approximately 10–15 μm, exhibited greater coactivity than distant spines (Extended Data Fig. 3a,b). Local coactivity increased during training, but neither baseline pairwise coactivity nor coactivity during reinforced events significantly predicted subsequent synaptic strengthening or weakening (Extended Data Fig. 3c-e). Thus, although neighboring synapses exhibited coordinated activity, the magnitude of their coactivity alone did not robustly explain subsequent plasticity.

We next asked whether the BMI task-relevant activity of spines provided a more informative predictor of synaptic plasticity than their general coactivity^25–27^. Because the BMI cursor was determined by the relative activity of E1 and E2, we examined whether E1 spine activity encoded cursor direction relative to the reinforced target (Fig. 3a). Cursor-related information was present even during early training (AUC = 0.63, p < 0.001), before reliable E1 control emerged (Fig. 3b), consistent with encoding of the experimentally defined task variable rather than a learned error representation. Notably, significant cursor-related information was recoverable from groups containing only a few spines (Fig. 3c). We refer to the joint activity configurations of identified spine groups used for decoding as synaptic codes.

**Figure 3.**
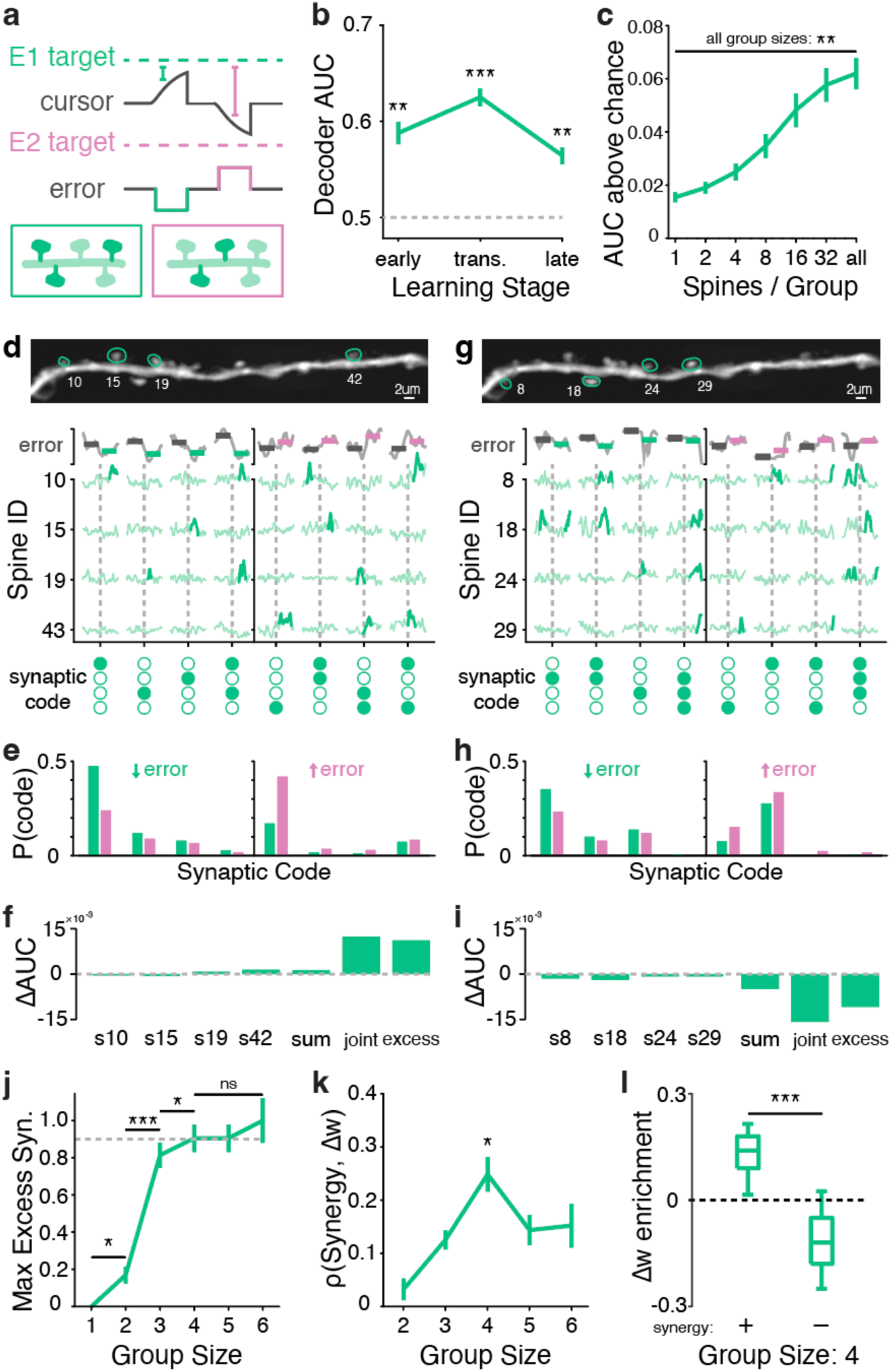
Plasticity is enriched in synergistic task-relevant synaptic groups. a, Schematic of the experimentally defined BMI cursor and target-related error, illustrating examples of synaptic codes used for decoding. b, Decoding performance for the direction of cursor error change (improving versus worsening) from E1 spine population activity across early, transition, and late training stages, quantified by area under the receiver operating characteristic curve (AUC). The dashed line indicates chance performance (AUC = 0.5). c, Decoding performance above chance as a function of the number of spines included in the decoder, illustrating task-related information recoverable from small synaptic groups. d–f, Representative positively synergistic synaptic group. d, Two-photon image identifying the constituent spines (top), corresponding spine calcium activity aligned with task-related error dynamics (middle), and examples of joint spine activity configurations (synaptic codes; bottom). e, Distribution of synaptic codes (as in d) during periods of improving (↓error) and worsening (↑error) cursor error. f, Contributions of the individual spines (from d), their summed contribution, and the joint group contribution to decoding performance. Positive excess denotes functional synergy. g–i, Corresponding analysis for a representative antagonistic synaptic group, illustrating its constituent spines (g), code distributions during improving and worsening cursor error (h), and negative excess decoding contribution relative to the sum of individual spine contributions (i). Scale bars in d,g, 2 μm. j, Maximum excess functional synergy, normalized to the largest measured value, as a function of synaptic group size. Excess synergy approached a plateau among small groups of approximately three to four spines. Dotted line denotes 90% of max. k, Correlation between early group functional synergy and subsequent synaptic plasticity polarity (fraction of strengthened - weakened spines) as a function of synaptic group size. l, Enrichment of synaptic plasticity polarity (LTP fraction - LTD fraction, relative to a shuffled null) among spines belonging to positively synergistic and antagonistic quadruplets identified during early training. Positive values indicate enrichment of strengthened spines; negative values indicate enrichment of weakened spines. Data are presented as mean ± s.e.m..

To determine whether these synaptic codes contributed independently or collectively to task-related information, we compared the decoding contribution of each synaptic group with the summed contributions of its individual members, defining the excess or deficit as functional synergy (Fig. 3d–i). This analysis revealed substantial heterogeneity: some groups exhibited positive synergy, contributing more to task decoding than expected from their individual members (Fig. 3d–f), whereas others exhibited negative synergy, or antagonism, contributing less than expected (Fig. 3g–i). The excess joint decoding contribution increased rapidly with group size, with triplets and quadruplets capturing approximately 80.2% and 89.6%, respectively, of the maximum observed excess contribution among up to six-spine groups (Fig. 3j). These values were not significantly different from those of larger groups, indicating that near-maximal joint decoding contributions could be achieved by small synaptic groups.

We next examined whether functional synergy measured during early training was associated with subsequent synaptic plasticity. Strikingly, early-stage group synergy was significantly correlated with subsequent synaptic plasticity, most prominently among spine quadruplets (ρ = 0.37, p = 0.032; Fig. 3k). Indeed, spines participating in positively synergistic groups during early training were enriched for strengthening by later training, whereas those participating in antagonistic groups were enriched for weakening (Fig. 3l).

This relationship with plasticity was strongest among groups exhibiting the largest positive and negative excess decoding contributions and persisted after accounting for baseline spine strength, event rate, temporal state, and separately for within-group activity correlations and task-prediction overlap (Extended Data Fig. 3f–h).

Finally, we examined whether functional synergy and the reinforcement-gated temporal plasticity rule (Fig. 2) captured overlapping or distinct relationships with subsequent synaptic changes. In spine-aware models incorporating both the baseline-state-by-reinforcement-timing interaction (S × R) and small-group functional context, positive synergy was associated with a bias in spine strengthening, whereas negative synergy was associated with a bias in spine weakening (Extended Data Fig. 3k). Together, these findings indicate that reinforcement-associated synaptic plasticity depends not only on when individual inputs are active, but also on how their collective activity relates to the reinforced task. Pre-existing temporal organization and task-related synaptic synergy thus provide complementary predictors of which inputs subsequently strengthen or weaken.

### Synaptic reweighting redistributes and temporally focuses synaptic ensemble dynamics

Having identified temporal and task-related features associated with subsequent synaptic strengthening and weakening, we next asked how this selective plasticity related to the emergence of organized synaptic activity during learning. The identities of spines recruited during E1 target hits became increasingly reproducible across training, with successive successful events exhibiting greater similarity in their constituent synaptic populations (Fig. 4a). Accordingly, recruitment became increasingly concentrated among a preferentially participating population of spines, which we term a probabilistic synaptic ensemble (Fig. 4b)^28,29^. This ensemble emerged progressively across learning, with substantial organization apparent during the behavioral transition and greater selectivity during late training. Importantly, the spines that ultimately dominated the learned ensemble were already active within the baseline and early learning input population (Extended Data Fig. 4a), suggesting that learning refined the recruitment of pre-existing synaptic identities rather than recruiting an entirely new population.

**Figure 4.**
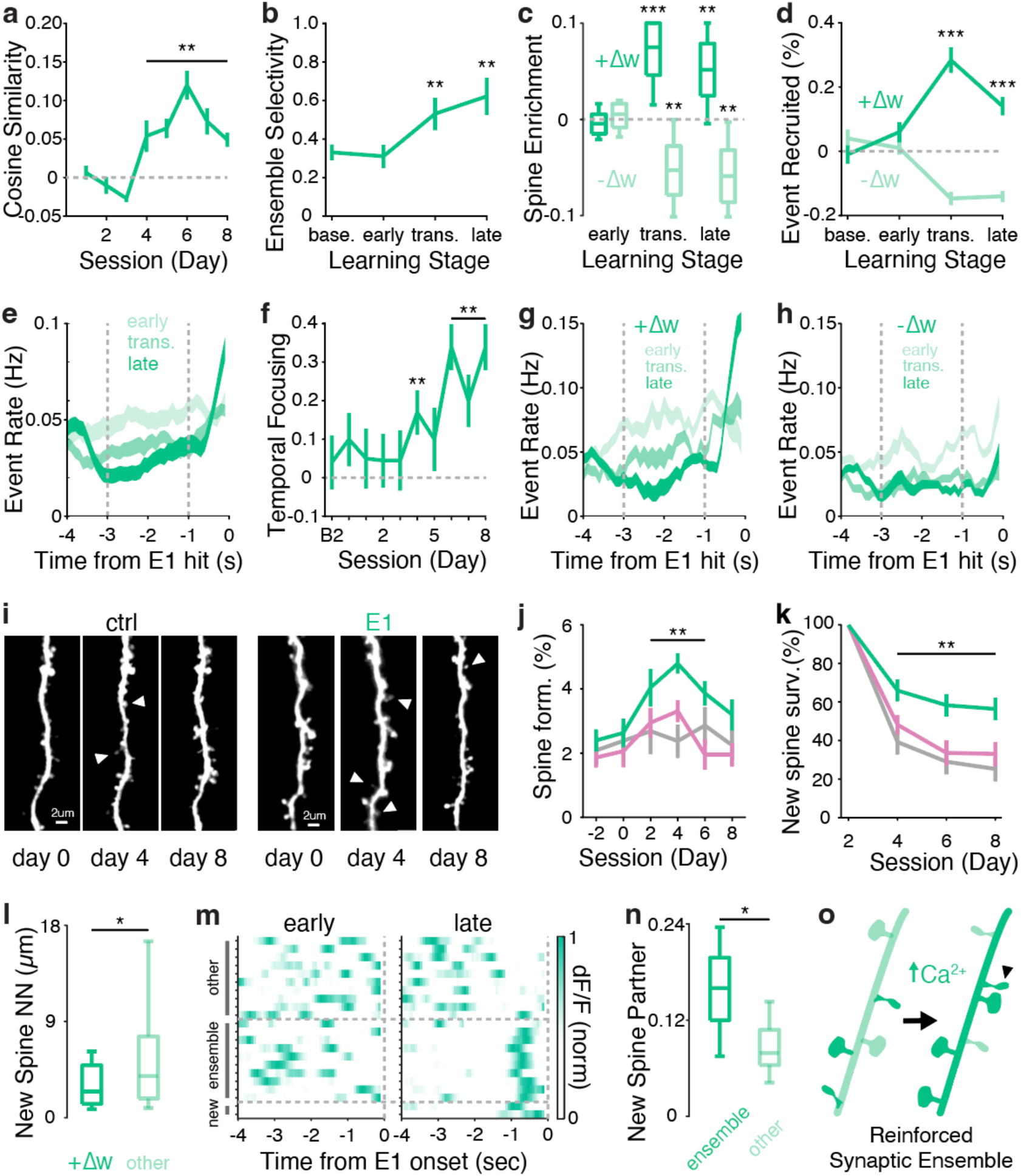
Synaptic reweighting redistributes and temporally focuses synaptic ensemble dynamics. a, Cosine similarity of synaptic identity recruitment during successful E1 events across training sessions, quantifying the increasing reproducibility of participating spine populations. b, Ensemble selectivity across baseline, early, transition and late training stages, quantifying the increasing concentration of recruitment among preferentially participating synaptic identities. c, Enrichment of strengthened (+Δw) and weakened (−Δw) spines within stage-defined synaptic ensembles across early, transition and late training. Positive values indicate enrichment; negative values indicate depletion relative to the corresponding expectation. d, Stage-dependent changes in event recruitment of strengthened and weakened spine populations during successful E1 events. e, Population-average spine calcium-event rates aligned to successful E1 dendritic events during early, transition and late training, illustrating progressive concentration of activity near dendritic-event onset. f, Temporal recruitment focusing across baseline and training sessions. g,h, Stage-resolved temporal recruitment profiles of strengthened (+Δw; g) and weakened (−Δw; h) spines preceding successful E1 dendritic events, illustrating their distinct contributions to temporal focusing. i, Representative longitudinal structural images of control and reinforced E1 dendritic segments at days 0, 4 and 8, illustrating newly formed spines (arrowheads). Scale bars, 2 μm. j, Spine formation across baseline and training sessions for the indicated populations. k, Survival of newly formed spines across training sessions for the indicated populations. l, Nearest-neighbor distance between newly formed spines and pre-existing strengthened (+Δw) or other spines. m, Representative activity maps of ensemble spines and newly formed spines during early and late training, aligned to successful E1 dendritic-event onset. n, Fraction of newly formed spine partners belonging to the learned synaptic ensemble compared with other synaptic partners. o, Schematic summarizing reinforcement-associated synaptic ensemble reorganization through selective enrichment of strengthened inputs, depletion of weakened inputs, temporal focusing of synaptic recruitment and incorporation of newly formed spines. Data are presented as mean ± s.e.m..

We next asked whether this selective recruitment was related to synaptic plasticity. Indeed, spines that underwent strengthening (+Δw) became progressively enriched within stage-specific synaptic ensembles, whereas weakened spines (−Δw) were progressively depleted from them (Fig. 4c). Consistent with this redistribution, the relative E1 hit recruitment of +Δw and -Δw spine populations diverged across learning, with strengthened inputs increasingly participating in E1 hits and weakened inputs decreasing in participation (Fig. 4d). These relationships remained evident after accounting for baseline differences in temporal state and recruitment probability (Extended Data Fig. 4b,c). Importantly, Δw was quantified from session-wide changes in isolated spine-event sizes, independently of these event-specific recruitment measurements. Thus, the enrichment of +Δw inputs and depletion of -Δw inputs among E1 hits reveal convergent changes in synaptic response strength and functional participation during reinforcement learning.

In addition to changes in ensemble composition, the temporal organization of spine calcium dynamics was progressively refined. During early training, spine activity preceding successful E1 events was distributed broadly across several seconds, whereas late training was characterized by increasingly concentrated activity immediately preceding dendritic-event onset (Fig. 4e). Quantification of this temporal recruitment revealed progressively stronger temporal focusing across sessions, particularly during late training (Fig. 4f). Notably, this reorganization occurred despite the absence of significant training-dependent changes in population-average isolated spine event rates or amplitudes (Fig. 2e,f), indicating a redistribution of synaptic activity in time without gross changes in synaptic activation.

We next examined how strengthened and weakened spines contributed to this temporal focusing. Both +Δw and -Δw spines exhibited reduced recruitment in the pre-dendritic interval as learning progressed (Fig. 4g,h). However, their activity diverged near dendritic-event onset: +Δw spines increasingly concentrated their recruitment within the final second preceding successful E1 events, whereas −Δw spines exhibited a broader reduction in recruitment without a comparable terminal concentration of activity. These distinct temporal profiles were most pronounced during late training and were robust to alternative temporal-window definitions (Extended Data Fig. 4d-f). Thus, the temporal focusing of the learned ensemble was associated with the selective preservation and concentration of strengthened inputs near successful dendritic events, alongside the broader withdrawal of weakened inputs.

Finally, we examined whether reinforcement learning also engaged synaptic rewiring of the E1 dendrite^30–34^. Our longitudinal structural imaging revealed the formation of new dendritic spines during training (Fig. 4i), with increased spine formation and enhanced stabilization on reinforced E1 dendrites (Fig. 4j,k). Among the tracked E1 spines across animals, 20 of 34 newly formed spines (58.8%) persisted through the final training session. New spines formed preferentially near pre-existing +Δw spines (Fig. 4l), suggesting that structural remodeling was spatially associated with dendritic regions undergoing synaptic strengthening^35–39^. Importantly, newly formed spines also exhibited preferential coactivation with members of the learned synaptic ensemble (Fig. 4m,n), supporting their functional incorporation into the selected input population. Together, these findings indicate that reinforcement learning refines a pre-existing synaptic population within a dendrite through the selective enrichment of strengthened inputs and depletion of weakened inputs, the temporal focusing of their recruitment during successful E1 events, and the incorporation of newly formed spines, resulting in a learned synaptic ensemble (Fig. 4o).

### Striatal dopamine receptor blockade impairs cortical synaptic reorganization

Corticospinal neurons send collaterals to the dorsal striatum, a critical component of the cortico-basal ganglia circuitry, where dopamine-dependent plasticity is implicated in reinforcement learning^40–45^. We therefore asked whether striatal dopamine receptor signaling contributes to the cortical synaptic plasticity and ensemble reorganization observed during VTA-mediated reinforcement.

To examine the contribution of striatal dopamine receptor signaling, we co-infused the D1 receptor (D1R) antagonist SCH23390 and D2 receptor (D2R) antagonist haloperidol into the dorsal striatum 10 minutes before BMI training and compared them with ACSF controls (Fig. 5a). Antagonist-treated animals displayed impaired acquisition of BMI control across sessions, although a measurable increase from early to late training remained (Fig. 5b). Because antagonist-treated animals achieved fewer E1 hits, they also received fewer optogenetic reinforcements: cumulative E1 reinforcements diverged between groups over training days (Fig. 5c), with antagonist-infused animals experiencing 31.2% fewer VTA stimulations than controls by the final training session. Impaired BMI performance was accompanied by reduced engagement of synaptic plasticity across training in antagonist-treated animals (Fig. 5d) and weaker synergy-associated plasticity (Fig. 5e), despite E1 control-related synergistic spine groups remaining detectable (Extended Data Fig. 5a-c). Notably, striatal receptor blockade did not produce gross changes in cortical dendritic or isolated spine calcium-event rates or amplitudes (Extended Data Fig. 5d,e).

**Figure 5.**
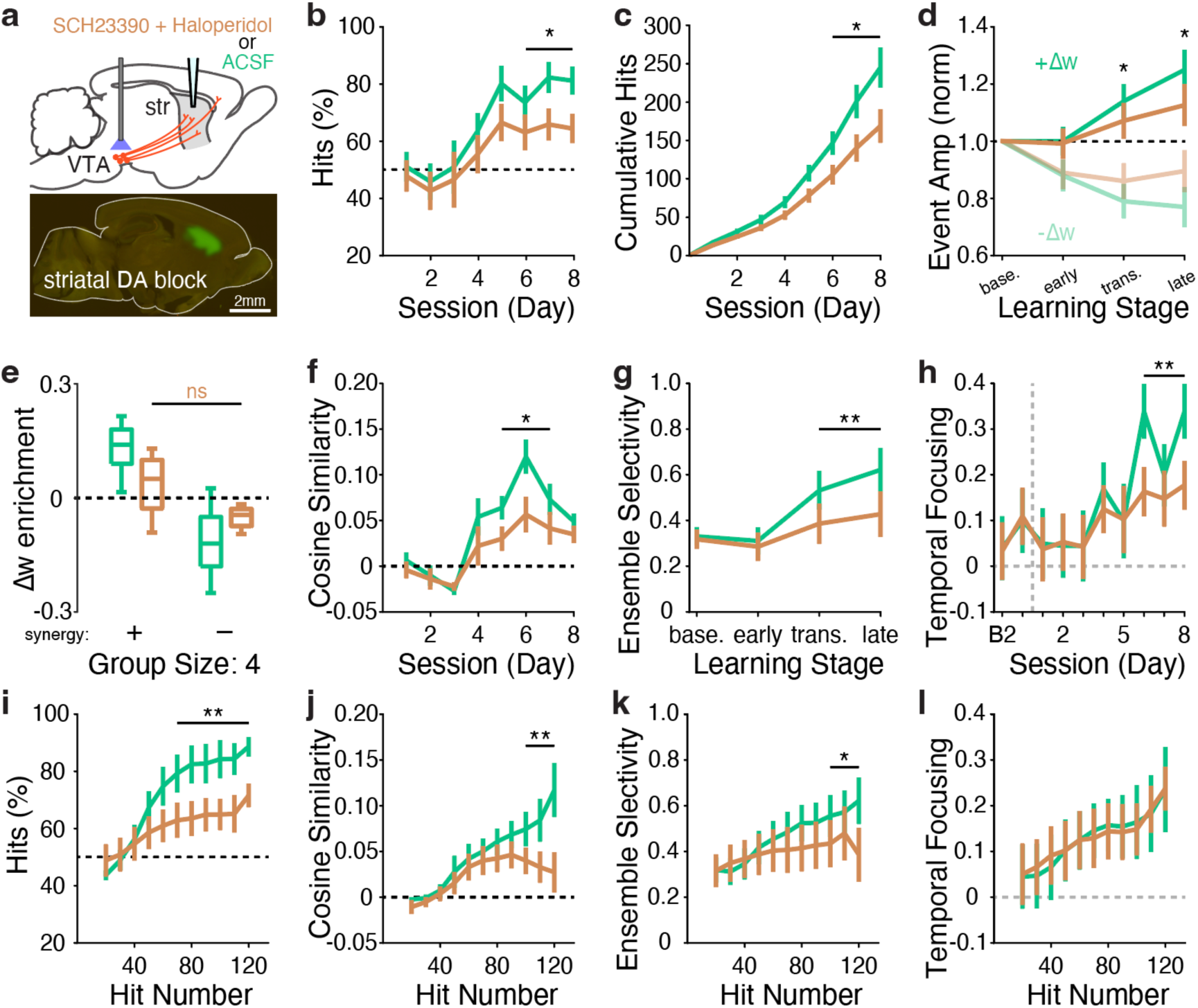
Striatal dopamine receptor blockade impairs cortical synaptic reorganization. a, Experimental design for unilateral local dorsal striatal infusion of the D1 receptor antagonist SCH23390 (10µM) and D2 receptor antagonist haloperidol (10µM) before optical BMI training, compared with artificial cerebrospinal fluid (ACSF) controls (note: these ACSF control animals were used for analyses in Figs. 1-4). Representative image illustrates the striatal infusion region. b, Fraction of successful E1 target hits across training sessions in ACSF- and antagonist-treated animals, illustrating decreased acquisition of E1-directed control following striatal dopamine receptor blockade. c, Cumulative E1 target hits, corresponding to cumulative optogenetic reinforcement events, across training sessions. d, Changes in normalized isolated spine calcium-event amplitude across baseline and training stages for strengthened (+Δw) and weakened (−Δw) populations under ACSF and antagonist treatment (comparing plasticity divergence; (+Δw) - (-Δw); across groups). e, Enrichment of synaptic plasticity (Δw) among constituent spines of positively synergistic and antagonistic quadruplets under ACSF and antagonist treatment (ns: non-significant). f, Cosine similarity of successful-event synaptic identity recruitment across training sessions, quantifying ensemble reproducibility. g, Ensemble selectivity across baseline, early, transition and late training stages. h, Temporal recruitment focusing across baseline and training sessions. i–l, Reinforcement-exposure-aligned analyses of BMI performance and synaptic ensemble organization. i, E1 target-hit percentage, j, ensemble reproducibility (cosine similarity), k, ensemble selectivity and l, temporal recruitment focusing as functions of cumulative E1 reinforcement count. Green, ACSF controls; brown, striatal D1R/D2R antagonist treatment. Data are presented as mean ± s.e.m..

Striatal dopamine receptor blockade was also associated with less pronounced reorganization of synaptic activity preceding E1 hits. Compared with controls, antagonist-treated animals showed reduced reactivation and concentration of spine activity into synaptic ensembles across learning (Fig. 5f,g), and significantly attenuated temporal focusing of spine dynamics (Fig. 5h). These results indicate that striatal receptor blockade disrupts reinforcement-driven cortical reorganization of synaptic activity.

We next asked whether the differences in BMI learning and cortical synaptic reorganization could be explained by differences in reinforcement exposure (Fig. 5b,c). We therefore compared learning and synaptic reorganization as a function of cumulative E1 reinforcement count rather than training session (Fig. 5i–l). At matched reinforcement exposure, antagonist-treated animals continued to show decreased task performance and attenuated ensemble refinement (Fig. 5i–k). Of note, striatal dopamine blockade did not completely abolish BMI learning or cortical synaptic reorganization, and temporal focusing also developed in both conditions, although the strongest session-dependent temporal focusing changes occurred late in training in controls, beyond the reinforcement counts reached by drug-infused animals (Fig. 5l). Thus, differences in cumulative reinforcement exposure alone did not account for the observed deficits in BMI learning and synaptic ensemble organization, indicating that normal striatal dopamine receptor signaling contributes to reinforcement-dependent reorganization of spines along specific cortical dendrites, beyond its effects on cumulative reinforcement exposure.

## Discussion

Using a closed-loop optical BMI where animals learn to control the activity of specific cortical dendrites and longitudinal imaging of identified dendritic spines in those dendrites, we unraveled several synaptic principles that underlie that learning. We found that synaptic plasticity was dependent not only on the timing of activity relative to reinforcement, but also on the pre-existing temporal organization of the activity of those synapses in relation to dendritic activity. Furthermore, plasticity was preferentially observed in specific spines that participated in small, synergistic groups with task-relevant activity. These selective changes were accompanied by the emergence of increasingly reproducible, temporally focused synaptic ensembles and by the incorporation of newly formed spines into these reinforced ensembles. Finally, striatal dopamine receptor blockade impaired both BMI learning and cortical synaptic reorganization. Together, these findings reveal how operant reinforcement of specific dendrites can be achieved by sculpting functional synaptic ensembles and establish optical BMIs as a powerful paradigm for investigating the synaptic rules of operant reinforcement learning.

A central implication of our findings concerns how synaptic eligibility is established during reinforcement learning. Classical three-factor learning rules propose that presynaptic activity, postsynaptic activity, and reinforcement interact to determine synaptic change^3,46,47^. Our findings suggest that reinforcement does not simply strengthen synapses that were recently active. Instead, whether an individual synapse strengthens or weakens is predicted by how the timing of its activity during reinforcement relates to its pre-existing temporal organization (Fig. 2). Thus, synapses are not equivalent when reinforcement occurs; their prior activity organization helps determine how subsequent recruitment relates to plasticity (Fig. 2n,o). This relationship was strongest during the transition to reliable E1 control, suggesting that temporal eligibility is particularly informative during acquisition. Together with previous demonstrations of temporally gated and compartment-specific plasticity^3,17,18,48,49^, our findings raise the possibility that ongoing dendritic activity establishes history-dependent eligibility states upon which reinforcement acts.

Synaptic selection was also associated with how the activity of groups of inputs collectively contributed to decoding task-relevant information. Although nearby spines exhibited coordinated activity, simple coactivity did not robustly explain their plasticity. Instead, functional synergy among small groups of spines predicted plasticity independently of the temporal plasticity rule (Fig. 3). Notably, these synaptic codes carried information about the experimentally defined cursor direction before reliable BMI control emerged, suggesting that task-related functional relationships were already present before learning. Together, these findings suggest that temporal eligibility and task-related synaptic interactions provide complementary constraints on synaptic selection, offering a potential framework for synaptic credit assignment^46,50,51^. Pre-existing synaptic organization may provide a repertoire of potential computations that reinforcement can selectively recruit and reshape. Notably, the observed functional synergy reflects nonadditive contributions to linear decoder performance, rather than formal information-theoretic synergy or direct evidence of nonlinear dendritic integration.

Our results also provide a synaptic ensemble perspective on the emergence of neuronal ensembles. Previous work has shown that cortical learning reorganizes population activity and functional connectivity^16,52,53^; our findings connect this reorganization to changes in the strength and recruitment of individual synaptic inputs. The convergence of independently measured synaptic strengthening and weakening with opposing changes in ensemble participation suggests that learning reshapes not only the responsiveness of individual inputs but also their probability of contributing to successful cortical activity. Importantly, increasingly reproducible E1 dendritic activation emerged from a probabilistic synaptic ensemble whose constituent combinations remained flexible across events. Such organization could support reliable cortical output despite variability in individual synaptic recruitment^54^. More broadly, differences in the pre-existing organization of synaptic inputs may influence which cortical neurons can most readily acquire particular reinforced activity patterns, potentially contributing to their selective recruitment into learned neuronal ensembles^10^. Together with the preferential incorporation of newly formed spines, these findings suggest that learning refines an existing functional population through selective reweighting, temporal focusing, and structural remodeling rather than establishing a rigid synaptic assembly.

The contribution of striatal dopamine receptor signaling places these findings within a broader framework of distributed reinforcement mechanisms. Dorsal striatal D1R/D2R blockade impaired BMI learning and associated cortical synaptic reorganization, with several deficits persisting after accounting for cumulative reinforcement exposure (Fig. 5). Nevertheless, residual learning and plasticity suggest that striatal dopamine signaling contributes to, but does not exclusively mediate, reinforcement-dependent cortical changes. The pathways linking optogenetic VTA stimulation to M1 synaptic plasticity remain unresolved and may involve cortico-basal ganglia-thalamocortical feedback^14,49,55,56^, direct cortical neuromodulation^57,58^, and/or parallel reinforcement-related signals^15,59,60^. Indeed, future experiments incorporating pathway-specific manipulations of excitatory, inhibitory^61–63^, and additional neuromodulatory circuits^64–67^ could establish how these signals interact to govern synaptic reinforcement.

Our measurements also define important boundaries for interpreting these findings. Spine-local calcium events provide an NMDAR-dependent readout of super-threshold synaptic activity, but do not directly capture subthreshold AMPAR-mediated events or local voltage dynamics, which also likely contribute to plasticity and learned dendritic control. Complementary iGluSnFR3 imaging did not reveal the strengthening-weakening divergence observed with GCaMP8m (Extended Data Fig. 2), highlighting the distinct aspects of synaptic function reported by extracellular glutamate and postsynaptic calcium signals. Furthermore, although dendritic calcium activity can reflect somatically coupled neuronal activity, dendrites can also support compartment-specific signaling and computations^29,68–74^. Our limited field of view prevented us from distinguishing these contributions or comparing plasticity across apical branches. Future extensions combining broader somatodendritic imaging with direct voltage measurements could determine how distinct dendritic activity states govern synaptic reinforcement. Lastly, whether these observed reinforcement-driven plasticity rules generalize beyond the apical dendrites of M1 layer 5 corticospinal neurons remains to be determined.

Together, our findings provide a synaptic-scale account of how reinforcement learning reorganizes cortical synaptic ensembles. Rather than uniformly strengthening active inputs, reinforcement acts within a pre-existing population of temporally organized and functionally interacting synapses. Individual temporal state and task-related synaptic synergy predict selective strengthening and weakening, while changes in synaptic recruitment, temporal organization, and structural composition accompany the emergence of increasingly reproducible learned ensembles. These findings suggest that reinforcement of specific dendritic activity sculpts synergistic task-relevant synaptic ensembles through selective plasticity and reorganization of small groups of inputs.

## METHODS

### Experimental Model and Subject Details

#### Mouse breeding and husbandry

Experiments were performed in DAT-FlpO (Jackson Laboratories, 040184) x Ai9 male and female at P60-80 (at the onset of the experiments). Mice were housed under 12hr:12hr light/dark cycle at an ambient temperature of 20-25°C and a relative humidity of 40-60%. All BMI imaging experiments were carried out at consistent times of day (7pm-10pm, every day) to avoid circadian and sleep related effects. All experiments and procedures were performed according to National Institutes of Health (NIH) guidelines, with lab protocols (AC-AABG2552, AC-AABG0559) approved by the Institutional Animal Care and Use Committee (IACUC) of Columbia University. Numbers of animals, dendrites and longitudinally tracked spines included in each analysis are reported in the corresponding figure legends and the reporting summary.

#### Viral expression and surgical procedures

To selectively express calcium and structural indicators in layer 5 corticospinal neurons of primary motor cortex (M1), we used DAT-FlpO mice crossed with Ai9 reporter mice. Viral constructs were acquired and/or generated in collaboration with the Columbia University Zuckerman Institute Molecular Tools Core Facility (RRID:SCR_026201). Mice received injections of a retrogradely transported Cre-expressing virus (pENN.AAV.hSyn.Cre.WPRE.hGH-retro; titer: 1.8E13 GC/mL; Addgene: 105553-AAVrg) into the cervical spinal cord, together with a Cre-dependent GCaMP8m virus (pGP-AAV-syn-FLEX-jGCaMP8m-WPRE; titer: 1.6E13 GC/mL; Addgene: 162378-AAV1) into the contralateral M1. This intersectional targeting strategy restricted GCaMP8m expression to corticospinal neurons projecting to the cervical spinal cord, enabling sparse labeling and longitudinal imaging of their apical dendrites and dendritic spines. To selectively express channelrhodopsin-2 (ChR2) in dopamine neurons of the ventral tegmental area (VTA), a Flp-dependent viral construct (pAAV-Ef1a-fDIO-hChR2(H134R)-EYFP; titer: 2.31E13 GC/mL; Addgene: 55639) was injected into the left VTA. Additional experiments involved co-injection of Cre-dependent iGluSnFR3 (pAAV-hsyn-FLEX-iGluSnFR3-v857-GPI; titer: 5.20E12 GC/mL; Addgene: 175181) and jRGECO1a (pAAV-hsyn-FLEX-jRGECO1a-deltaNES; titer: 6.80E13 GC/mL; Addgene: 100853) into M1 DAT-FlpO mice (Extended Data Fig. 2).

Surgeries were performed under sterile conditions and isoflurane anesthesia (1%-5%, plus oxygen at 1-1.5 l/min) using a stereotaxic apparatus (Kopf Instruments). Preoperative analgesia consisted of subcutaneous carprofen (5 mg/kg) or sustained-release buprenorphine (0.5–1 mg/kg), together with bupivacaine (2 mg/kg). For cortical viral injections, a midline incision was made to expose the skull, and a small craniotomy was performed over the left M1. 100nL of the Cre-dependent GCaMP8m virus was injected at two sites in forelimb motor cortex, located 0.5 and 1.0 mm AP, 1.5 mm ML, and 700 μm DV (below the pial surface). Viral injections were performed using a pulled glass micropipette (Nanoject III injector; Drummond Scientific, USA) that was maintained in place for 10 mins before withdrawal to minimize viral reflux. Flp-dependent ChR2 was injected into the left VTA at a 30° angle at -3.52 mm AP, 0.35 mm ML, and 4.3 mm DV.

For spinal cord injections, a midline incision was made to expose the cervical spinal column, and the overlying musculature was carefully retracted. The spinal column was stabilized using spinal clamps to minimize movement during injections. Following exposure of the spinal cord, a pulled glass micropipette was used to deliver retrograde Cre virus into the cervical spinal gray matter. Injections targeted spinal segments C3–C7, with 20 penetration(s) per segment (10 nL at 0.1 mm DV increments) for a total volume of 200 nL delivered at each location. Following injections, the incision was sutured closed, and mice were monitored during postoperative recovery.

For chronic two-photon imaging, mice were implanted with a 5mm cranial window over left M1 as previously described^32^. An optical fiber (200/230 µm diameter, 6 mm length; Plexon) was implanted above the left VTA for optogenetic stimulation.

### Histological validation

For histological validation of VTA-targeting and VTA-ChR2 expression (Extended Data Fig. 1) and validation of dorsal striatal infusion sites (Extended Data Fig. 5), mice were deeply anesthetized with isoflurane and transcardially perfused with PBS, followed by ice-cold 4% paraformaldehyde. Brains were then removed for histological analysis. Coronal or sagittal sections were cut at 60 µm using a Leica VT1000 vibratome. The tissue was rinsed twice in 1x PBS and then DAPI (Sigma D9542; 1:1000) was used as a counterstain in all experiments. Coronal or sagittal sections were serially mounted on slides and sections were imaged using an automated slide scanner (Nikon AZ100 Multizoom microscope) equipped with a 4x 0.4NA Plan Apo objective (Nikon Instruments Inc) and P200 slide loader (Prior Scientific), controlled by NIS-Elements using custom acquisition scripts (Nikon Instruments Inc.).

### Two-photon calcium imaging

All imaging experiments were performed using a Bergamo II two-photon laser-scanning microscope (Thorlabs, Inc.; Columbia University Zuckerman Institute Cellular Imaging Core Facility, RRID:SCR_027061) equipped with 8-kHz resonant-galvanometer scanning mirrors and controlled using ThorImage software. Functional imaging was performed at 30 frames per second with a resolution of 512×1024 pixels or 512×1024 pixels, corresponding to a field of view of approximately 52×52 um or 52×104 µm (or 104 × 208 μm depending on the optical zoom factor). Fluorescence was collected using two amplified, non-cooled gallium arsenide phosphide (GaAsP) photomultiplier tubes (PMTs). Emitted fluorescence was separated into green and red channels using a 565-nm long-pass dichroic mirror and subsequently filtered using 525/39-nm and 593/40-nm bandpass filters, respectively.

Imaging was performed using a Nikon 16x, 0.8 NA water-immersion objective. A tunable titanium–sapphire laser (Coherent Chameleon Vision S) was used for two-photon excitation at 920 nm, with a pulse duration of approximately 75 fs and an 80-MHz repetition rate. Laser dispersion compensation was adjusted to maximize fluorescence intensity, and excitation power was regulated using a Pockels cell to maintain fluorescence measurements within the linear detection range. Laser power at the objective was maintained at <40 mW during functional imaging.

GCaMP8m and tdTomato fluorescence were simultaneously recorded from the apical dendrites of sparsely labeled layer 5 corticospinal neurons in M1. Imaging fields were selected to include dendritic segments corresponding to the E1 and E2 populations used for optical BMI control. The same identified dendritic segments and their associated spines were longitudinally imaged across two baseline sessions (B2 and B1) and eight training sessions (T1–T8). Dendritic morphology, spine positions, and local anatomical landmarks were used to identify corresponding structures across imaging sessions.

For longitudinal structural measurements, additional high-resolution images were acquired at 1040 nm. These images were used to identify persistent, newly formed, and eliminated dendritic spines across training.

### Closed-loop Optical Brain-Machine Interface

#### Experimental design

We developed a closed-loop optical brain–machine interface (BMI) in which mice learned to control the activity of specific cortical dendrites to obtain optogenetic reinforcement. Two predefined apical dendritic regions of interest (ROIs), designated E1 and E2, served as the online BMI control signals. Each ROI corresponded to a single apical dendritic branch of a layer 5 corticospinal neuron in primary motor cortex (M1). Mice received continuous auditory feedback corresponding to the difference between the normalized activity of the E1 and E2 dendrites. Increased E1 activity relative to E2 moved the cursor in the positive direction toward a high-frequency, reinforced target, whereas increased E2 activity relative to E1 moved the cursor in the negative direction toward a low-frequency, nonreinforced target.

Target acquisition was determined by thresholds established during baseline calibration. Acquisition of the E1 target triggered optogenetic stimulation of the ventral tegmental area (VTA) dopamine neurons, whereas acquisition of the E2 target did not result in reinforcement. Optical stimulation was delivered unilaterally using a 473-nm laser coupled to the implanted optical fiber through a patch cord. Each reinforcement consisted of a 600-ms stimulation train containing 15 light pulses (10-ms pulse width, 25 Hz; 25% duty cycle). Laser power was calibrated before each training session to approximately 10 mW at the fiber tip using a reference fiber of comparable optical transmission efficiency. Following target acquisition, a 2-s intertrial interval was imposed. Training sessions lasted a maximum of 20 min or until 60 optogenetic reinforcement events had been delivered.

#### Online fluorescence processing and BMI calibration

Custom MATLAB software accessed the microscope’s raw imaging files during acquisition and extracted fluorescence signals from predefined dendritic ROIs in real time. Fluorescence measurements were updated at the imaging frame rate (30 Hz). The first 100 frames of each recording were excluded from online fluorescence processing. For each ROI, fluorescence was smoothed using a causal four-frame moving average, corresponding to approximately 133 ms. Baseline fluorescence ((F_0)) was initialized during calibration and subsequently calculated using a rolling 120-s mean of the fluorescence signal. Normalized calcium activity was calculated as: ΔF/F_0_ = (F-F_0_)/F_0_.

Before BMI training, baseline fluorescence activity was collected to calibrate the decoder and establish target thresholds. To account for differences in fluorescence amplitude between the two dendritic ROIs, each signal was independently range-normalized using calibration-derived minimum and maximum values. Target thresholds were established from the distribution of spontaneous cursor activity, using an iterative percentile-based calibration procedure designed to yield approximately one target acquisition per 60 s. The target-search algorithm evaluated percentiles between 50 and 100, with a maximum of 100 iterations. Calibration parameters were subsequently used to define the target boundaries during BMI training.

#### Online BMI decoder and auditory feedback

The BMI control signal was calculated using a linear decoder that combined the normalized activity of the two dendritic ROIs. Each ROI was assigned to one of two ensembles (E1 and E2), and the corresponding ensemble projection vectors were normalized to unit Euclidean length. The difference between the two projection vectors was subsequently normalized, producing a decoder that weighted the E1 and E2 signals equally and oppositely.

For consistency, the BMI cursor was defined using the E1-positive convention:

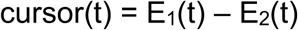

where E_1_(t) and E_2_(t) represent the independently normalized calcium activity of the corresponding dendritic ROIs. Thus, positive cursor values indicated preferential E1 activity and movement toward the reinforced target, whereas negative values indicated preferential E2 activity and movement toward the nonreinforced target. For single-ROI ensembles, the normalized decoder output was proportional to this difference, with a common scaling factor determined by the decoder normalization.

Cursor position was continuously converted into auditory feedback spanning 6–19 kHz using a calibrated piecewise exponential mapping. The central cursor position was defined as the median of the spontaneous cursor distribution recorded during the pretraining calibration period (50th percentile), rather than the midpoint between the E1 and E2 target thresholds. A central region, bounded by the 25th and 75th percentiles of the calibration distribution, was additionally defined to determine when the cursor had returned to its central state before initiation of a subsequent trial. The central cursor position was mapped to approximately 10.7 kHz, corresponding to the geometric midpoint of the auditory frequency range, whereas the E1 and E2 target thresholds were mapped to 19 and 6 kHz, respectively. Separate exponential mappings were applied on either side of the center, providing continuous auditory feedback proportional to deviations from the calibrated spontaneous cursor state.

#### Experimental timeline

Mice underwent two pretraining baseline imaging sessions (B2 and B1), followed by eight consecutive daily BMI training sessions (T1–T8). During baseline sessions, dendritic and spine calcium activity was recorded in the absence of contingent optogenetic reinforcement. During training, mice controlled the auditory cursor through E1 and E2 dendritic activity, with E1 target acquisition triggering VTA dopamine neuron stimulation.

For descriptive analyses, training sessions were grouped into early (T1–T3), transition (T4– T5), and late (T6–T8) stages, reflecting the temporal progression of BMI learning. These stages were used to examine how dendritic activity, synaptic plasticity, and synaptic ensemble organization evolved before, during, and after the emergence of reliable E1-directed control.

#### BMI performance analysis

BMI performance was quantified from the recorded timestamps of trial initiation, E1 and E2 target acquisition, and the corresponding dendritic activity. Behavioral events were synchronized to two-photon imaging frames using the recorded acquisition and synchronization signals. Target acquisitions were identified from the experimental event timestamps rather than inferred from detected dendritic calcium events.

For each training session, the number of E1 and E2 target acquisitions was quantified separately. Relative performance was expressed as the percentage of all target acquisitions directed toward E1:

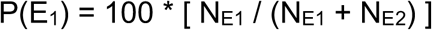

where N_E1_ and N_E2_ represent the numbers of E1 and E2 target acquisitions, respectively. An E1 hit fraction of 50% indicated equal acquisition of the two targets, whereas values greater than 50% indicated preferential acquisition of the reinforced E1 target. Target acquisition rates were additionally calculated as the number of acquisitions per minute of valid recording time (Extended Fig. 1).

To assess changes in auditory cursor occupancy across training, E1 and E2 fluorescence signals were reconstructed using a fixed, animal-specific calibration derived from the two baseline imaging sessions (B1 and B2). For each baseline session, calibration parameters were estimated independently from the recorded dendritic activity and included the normalization offsets and ranges for the E1 and E2 signals, the central cursor position, and the E1 and E2 target thresholds. These parameters were then averaged across the two baseline sessions and held constant when reconstructing cursor trajectories and auditory frequencies throughout training (T1–T8). This ensured that changes in cursor occupancy were measured relative to the same baseline-defined cursor coordinates, rather than being influenced by session-specific recalibration.

Cursor occupancy was quantified as the fraction of valid trial time spent within defined auditory-frequency ranges. E1-target occupancy gain was calculated as the fraction of time spent at the reinforced E1 target (19 kHz) during each training session divided by the corresponding fraction during the first training session (T1). High-versus-low cursor occupancy gain was calculated by first determining the fraction of extreme-frequency occupancy directed toward E1, defined as occupancy in the high-frequency range (16–19 kHz) divided by the combined occupancy in the high- and low-frequency ranges (16–19 kHz and 6–9 kHz, respectively). This fraction was then divided by its corresponding value during T1. Both measures were calculated independently for each animal, such that a value of 1 indicated no change relative to T1.

To quantify the emergence of stereotyped dendritic control, individual cursor trajectories were reconstructed from trial initiation to target acquisition and classified according to their recorded E1 or E2 outcomes. Trajectory similarity was evaluated within and across training sessions after normalization of trial duration and cursor position, as detailed below. These analyses assessed the reproducibility of cursor dynamics associated with reinforced E1-directed and nonreinforced E2-directed control.

#### Pharmacological manipulations

To investigate the contribution of striatal dopamine receptor signaling to cortical reinforcement learning, mice received local infusions of the D1 dopamine receptor antagonist SCH23390 (10 μM; Sigma: D054) and the D2 dopamine receptor antagonist haloperidol (10 μM; Sigma: H1512) into the dorsal striatum ipsilateral to the imaged M1 hemisphere. Antagonists were co-administered 10 min before each BMI training session. Control mice (same as Figs. 1-4) received equivalent infusions of artificial cerebrospinal fluid (ACSF) containing 125 mM NaCl, 2.5 mM KCl, 1.25 mM NaH2PO4, 25 mM NaHCO3, 15 mM glucose, 2 mM CaCl2 and 1 mM MgCl2, oxygenated with 95% O2 and 5% CO2 (300–305 mOsm, pH 7.4).

Infusions were performed through a silicone access port incorporated into the chronic cranial window. A glass micropipette connected to a hydraulic micromanipulator (Nanoject III Injector) was inserted through the access port and advanced at an angle through the cortex to reach the dorsal striatum. The injection site was targeted at approximately +0.5 mm AP, 2.65 mm ML, and 3.5 mm DV. Dopamine receptor antagonists were prepared in ACSF. The same procedure was followed for NMDA receptor antagonist (R-CPP, 10 µM; Sigma: 504839) infusions into local M1 (Extended Data Fig. 2).

### Image Processing and Event Extraction

#### Alignment of imaging, behavior, and reinforcement

Behavioral-event times and imaging acquisition were aligned using Thorsync-triggered signals for imaging frames, trial initiation, E1 and E2 targets, optogenetic triggers and shutter state (acquired at 30kHz). Digital traces were thresholded and their rising edges converted to imaging-frame indices. The resulting session records retained the precise recorded frames of target acquisition, trial initiation and stimulation. Analyses distinguished BMI target hits from detected dendritic calcium events: the former were recorded task outcomes, whereas the latter were extracted independently from fluorescence recordings.

#### Image registration and ROI extraction

Raw imaging data were separated into green and red channels where the acquisition stored interleaved frames. Movies were registered with nonrigid NoRMCorre using a common reference template and batch processing. Shutter-closed frames were excluded from template selection and were replaced temporarily by neighboring valid frames for the purposes of motion estimation. Dendrite and spine ROIs were manually identified on registered projection images.

#### Fluorescence correction

For each ROI, the extracted green fluorescence trace was detrended and corrected for fluorescence fluctuations shared with the red structural channel. Spine-local activity was further estimated by subtracting the component attributable to the parent dendrite, generating a dendrite-subtracted green signal (G_ds_). Both red-corrected and dendrite-subtracted traces were retained, allowing event metrics to be checked against their non-subtracted counterparts.

#### Calcium-event detection and classification

Dendritic and spine calcium transients were detected from processed fluorescence traces using custom MATLAB routines. Candidate events were identified as periods in which fluorescence exceeded a threshold of three times the estimated baseline noise, with baseline fluorescence estimated from the median signal and noise estimated using the median absolute deviation (MAD; σ = 1.4826 × MAD). Candidate events were required to remain above threshold for at least 100 ms, and events separated by intervals of 100 ms or less were merged. Events were additionally required to reach a minimum peak amplitude of three times the estimated baseline noise. Event onset times were refined by searching backward from the initial threshold crossing to the point at which fluorescence fell below one standard deviation above baseline, within a maximum preceding window of 1 s. Event onset, peak, and offset times were subsequently used to quantify event frequency, amplitude, duration, and temporal relationships between individual spines and their parent dendrites.

To distinguish spine-local calcium activity from signals associated with parent-dendrite activation, detected spine events were classified according to their temporal relationship with dendritic calcium events. Spine events were classified as isolated if they did not overlap a dendritic event or if their onset preceded that of an overlapping dendritic event. Events beginning during an overlapping dendritic event were classified as coincident. For analyses of functional synaptic strength and plasticity, we exclusively used isolated spine events, thereby minimizing contributions from dendrite-wide calcium activity to measurements of individual spine-event amplitude and area. Analyses of synaptic recruitment timing, coactivity, and ensemble organization used the corresponding detected spine-event populations, with additional event-exclusion criteria applied where indicated. The robustness of event detection was additionally evaluated by varying the detection thresholds and temporal criteria.

#### Across-session BMI and activity analyses

Event rates were calculated as the number of detected events divided by valid, non-padded recording duration. Dendritic and spine activity metrics were summarized within imaging session and across training days. Where baseline normalization was used, each animal’s E1 and E2 measurements were divided by the corresponding role-specific median of the B2 and B1 sessions. Target-hit fractions and rates were calculated from the actual recorded E1 and E2 target events. For trajectory analyses, cursor trajectories were reconstructed using the E1-positive cursor signal derived from dendritic fluorescence, with calibration parameters established from the two baseline sessions (B2 and B1). Trajectories were separated according to E1 and E2 target-acquisition outcomes. To account for differences in trial duration, each trajectory was temporally normalized by assigning trial initiation and target acquisition to normalized times of 0 and 1, respectively, and linearly interpolating the intervening cursor values onto a common grid of 100 equally spaced time points. This procedure preserved the amplitude and shape of individual cursor trajectories while allowing direct comparisons of their relative temporal evolution across trials of different durations.

Within-session trajectory stereotypy was quantified as the mean pairwise Pearson correlation coefficient between time-normalized cursor trajectories. Correlations were calculated between all unique pairs of trajectories within each session and outcome category. Individual correlation coefficients were Fisher z-transformed before averaging and subsequently back-transformed to obtain the session-level stereotypy measure. Higher values indicated greater trial-to-trial reproducibility of cursor trajectories.

Across-session trajectory stereotypy was quantified between consecutive training sessions using the same baseline-calibrated, time-normalized cursor trajectories. For each pair of consecutive sessions, Pearson correlation coefficients were calculated between all valid pairs of trajectories from the preceding and current sessions, separately for E1 and E2 target-acquisition outcomes. Correlation coefficients were Fisher z-transformed before averaging and subsequently back-transformed to obtain a single across-session stereotypy measure for each outcome and session transition. Higher values indicated greater similarity of cursor trajectories across successive training days.

To assess the robustness of these findings to differences in cursor amplitude and target acquisition, additional analyses used trajectories normalized to their starting and terminal cursor positions. These endpoint-normalized analyses emphasized trajectory shape independently of overall cursor displacement.

### Longitudinal Synaptic Plasticity

#### Functional spine strength and plasticity

For each spine and session, peak amplitudes and integrated areas of dendrite-subtracted calcium events were summarized separately across isolated, coincident and all detected events. Functional synaptic strength was quantified primarily from isolated spine calcium events, minimizing contributions from shared dendritic fluorescence. Four measurements were extracted for each spine: mean event amplitude, median event amplitude, mean event area and median event area.

For each measurement, baseline strength was calculated by averaging values from the two baseline sessions (B1 and B2), and late-training strength was calculated by averaging values from T7 and T8. A composite logarithmic plasticity measure was then defined as the median log₂ fold change across the four measurements:

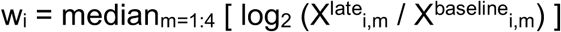

where X_i,m_ denotes measurement (m) for spine (i).

For categorical analyses separating strengthened, stable and weakened spines, the composite log₂ fold change was converted into percentage plasticity as:

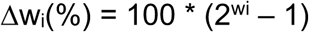

Spines exhibiting greater than 10% strengthening were classified as strengthened (+Δw), those exhibiting greater than 10% weakening were classified as weakened (−Δw), and those exhibiting changes between −10% and +10% were classified as stable. These classifications were used for categorical visualization and group comparisons, with additional sensitivity analyses using alternative thresholds and thresholds derived from baseline variability.

For analyses of the reinforcement-dependent temporal plasticity rule, the primary continuous plasticity endpoint was a baseline-adjusted late strength-state residual. Initial and late strength states were calculated by converting each of the four isolated-event measurements into centered percentile scores across spines and averaging the resulting scores. The late strength-state residual was then obtained after linearly accounting for the relationship between initial and late strength states. This provided a continuous measure of subsequent synaptic plasticity that accounted for differences in initial functional strength.

#### Pre-existing synaptic temporal organization parameter S

To characterize the pre-existing temporal organization of synaptic activity, isolated spine calcium-event onsets were aligned to subsequent parent-dendritic calcium-event onsets during the two baseline sessions (B2 and B1). For each spine, the distribution of pre-dendritic recruitment times was divided into eight equal temporal bins spanning approximately 3.6 s before dendritic-event onset. These bins were grouped into three contiguous sectors: distal (approximately −3.60 to −2.25 s), intermediate (−2.25 to −1.35 s), and proximal (−1.35 to 0 s).

The primary baseline temporal-state measure, S_i_ (where *i* is a unique spine), was defined as the difference between the probabilities of spine recruitment within the proximal and distal sectors:

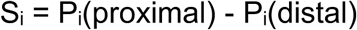

Positive values indicated preferential recruitment closer to dendritic-event onset, whereas negative values indicated preferential recruitment during earlier portions of the pre-event interval. The intermediate sector contributed to the underlying temporal distribution but was excluded from the sector contrast. S_i_ was treated as a continuous predictor of subsequent synaptic plasticity rather than evidence of discrete synaptic classes.

#### Reinforcement-associated recruitment timing and temporal plasticity rule

During BMI training, spine-event onsets were aligned to E1 dendritic events preceding successful E1 target acquisition. The recruitment-timing variable R summarized the difference between late/proximal and early/distal timing occupancy in the relevant reinforced-event context. Importantly, R was defined with respect to the dendritic-event onset, not as a direct measurement of the delay to the VTA stimulation pulse. Reinforced and nonreinforced event contexts and early, transition and late training stages were retained separately.

To assess whether reinforcement-associated synaptic plasticity depended on the interaction between baseline synaptic timing and reinforcement timing, we fitted a multiple linear regression model:

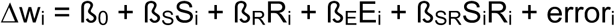

where Δw_i_ denotes the baseline-adjusted change in functional synaptic strength of spine (i), S_i_ represents its baseline temporal bias, and R_i_ represents its reinforcement-associated temporal bias. Total reinforcement-associated exposure E_i_ was calculated by summing spine-event exposure across the proximal and distal temporal sectors. Including total exposure as a covariate allowed us to assess whether the interaction between baseline timing and reinforcement-associated timing predicted synaptic plasticity independently of the overall amount of reinforcement-associated spine activity. The interaction coefficient ß_SR_ quantified this relationship after accounting for the individual contributions of baseline temporal bias, reinforcement-associated temporal bias, and total exposure.

### Spine coactivity and task-relevant synergy

#### Spatial distance and local spine coactivity

Interspine distances were calculated as Euclidean path-distances (along the dendrite) between ROI centroids, converted from pixels to micrometers using 0.102 µm / pixel. Spine-event rasters were binarized over the duration of each detected event. For pairs of spines, observed coactivity was measured as the number of frames during which both spines were active. Chance coactivity was estimated from independently permuted or shifted binary traces, and chance-subtracted activity was calculated by subtracting the median shuffled estimate. Some analyses restricted coactivity to local neighbors within a 10-micrometer radius. To distinguish spine coactivity from shared dendritic calcium activity, the dendrite-clean coactivity analysis excluded spine events overlapping a parent-dendrite exclusion interval extending approximately 1 s on either side of the dendritic event.

#### Decoding task-related information from spine activity

To quantify synaptic information about task performance, we trained ridge-regularized linear decoders to distinguish epochs in which cursor error was improving from those in which it was worsening. Cursor error was defined relative to the reinforced E1 target, and its temporal derivative was calculated from the reconstructed auditory-feedback cursor. Continuous activity was divided into non-overlapping 1-s epochs, and changes in cursor error were evaluated over a corresponding 1-s window. Epochs in the lower and upper quartiles of the cursor-error derivative distribution were classified as improving and worsening, respectively; intermediate epochs were excluded from binary classification. The 2-s period following target acquisition was excluded to minimize contributions from reinforcement and trial-reset transients.

For each spine, predictors consisted of continuous-valued estimates of calcium-event onset rate averaged within each epoch. To isolate information carried by the configuration of synaptic activity beyond differences in overall activity and task state, spine predictors were adjusted for mean population spine-event rate, current absolute cursor error, and the number of dendritic calcium events occurring within ±0.5 s of the epoch timestamp. Confound regression was performed using the training data within each cross-validation fold, and the fitted transformations were subsequently applied to held-out data.

Linear decoders were fitted using ridge-regularized least squares with a regularization parameter of λ = 10. Predictors were standardized using training-fold statistics, and inverse-frequency observation weights were applied to balance the contributions of improving and worsening epochs. Decoder performance was assessed using leave-one-session-out cross-validation, with discrimination quantified by the area under the receiver operating characteristic curve (ROC AUC; chance = 0.5). Analyses were performed separately for early (T1–T3), transition (T4–T5), and late (T6–T8) training stages.

To assess how task-related information depended on the number and identity of contributing spines, decoder performance was compared across spine subsets of different sizes. Individual spine contributions were quantified as the reduction in cross-validated decoder performance following removal of that spine. The joint contribution of a group of spines was similarly quantified by removing all members of the group. Excess synergy was defined as the joint reduction in decoder performance minus the sum of individual spine contributions, with positive and negative values indicating supra-additive and sub-additive contributions to decoding, respectively.

Statistical significance of decoding performance was assessed using permutation-based null distributions generated by shuffling improving and worsening labels and repeating model fitting and cross-validation. We used 1,000 permutations, with two-sided significance calculated by comparing the absolute deviation of observed ROC AUC from chance (0.5) against the corresponding null distribution.

#### Individual-spine contributions and functional group synergy

The contribution of individual spines and groups of spines to task-related information was quantified using leave-feature-out analyses of the ridge-regularized decoder described above. For a group of spines *G*, the joint informational contribution was defined as the reduction in cross-validated decoder performance following removal of all group members:

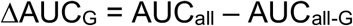

The contribution of each spine i was defined analogously as the difference between full-model AUC and AUC after removal of that spine. Group excess synergy was calculated as:

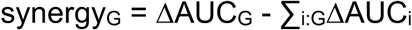

Positive values indicated that the joint contribution of a spine group exceeded the sum of its individual contributions, whereas negative values indicated subadditive or antagonistic contributions. This operational measure quantified nonadditivity in the contribution of synaptic activity patterns to the performance of a regularized linear decoder, rather than synergy as formally defined in information theory, and did not imply nonlinear dendritic integration. Excess synergy was examined across groups of two to six spines. Candidate groups were assembled from the longitudinally identified spine population, with all combinations evaluated when computationally feasible and otherwise sampled without replacement from the space of possible combinations. Group contributions were calculated using the same cross-validation framework and confound-control procedures as the full decoder.

To determine the characteristic size of functionally synergistic spine groups, maximum positive excess decoding contribution was calculated separately for each group size and normalized to the largest value observed across the tested sizes. The resulting normalized synergy curve was used to determine the smallest group size approaching 90% of the maximum measured synergy. Individual spine groups were also ranked by excess synergy to identify strongly synergistic and antagonistic configurations for subsequent analyses.

#### Controls for correlated activity and redundancy contributing to decoding

To assess whether excess group decoding contributions could arise from correlations or redundancy among synaptic predictors, we performed additional analyses using the same early-training spine event-rate representations, decoder configuration, and original spine-group memberships. For each group, we quantified the mean absolute pairwise correlation among constituent spine activity rates (within-group) and between group members and spines outside the group (outside-group). We then examined how these measures related to the joint group decoding contribution, the summed individual-spine contributions, and their difference (excess ΔAUC). Within-group activity correlations were generally weak and showed little association with excess decoding contributions (quadruplets: ρ = −0.019). Outside-group correlations were more strongly associated with excess (ρ = 0.310), but were positively, rather than negatively, associated with joint decoding contributions (ρ = 0.278), inconsistent with a simple compensation mechanism whereby correlated outside-group spines substitute for the removed group.

We further assessed task-related overlap among group members by training individual-spine decoders using the same leave-one-session-out cross-validation procedure. Similarity between their continuous held-out decoder scores was quantified using mean absolute pairwise correlations, including correlations calculated separately within improving and worsening cursor-error epochs. Task-score overlap showed a modest positive association with excess decoding contributions (quadruplets: ρ = 0.194). To assess whether this overlap accounted for the relationship between early excess decoding contribution and later plasticity, we calculated rank-based partial correlations after adjusting both variables for within-group task-score overlap. The association between excess decoding contribution and subsequent plasticity remained largely unchanged (quadruplets: original ρ = 0.373; adjusted ρ = 0.365, spine-identity permutation p = 0.044). Permutations shuffled future plasticity labels across eligible spine identities while preserving the original group memberships and overlap structure. In contrast, adjustment for outside-group activity correlations reduced the association (adjusted ρ = 0.271, p = 0.157), suggesting that broader population coupling may contribute to the observed relationship without necessarily reflecting redundant task information.

Finally, we examined progressive group-member ablations, including all nonempty subsets of the sampled groups, to characterize how decoder performance changed as additional constituent spines were removed. High-excess groups exhibited relatively small individual-spine decoding losses but progressively greater losses with joint removal, consistent with partially substitutable contributions. However, this pattern is partly inherent to the definition of excess ΔAUC and therefore does not independently establish redundancy. Together, these analyses indicate that simple within-group activity collinearity and measured task-prediction overlap do not fully account for the relationship between functional synergy and subsequent plasticity. Nevertheless, correlated predictors may contribute to decoder nonadditivity, and the operational excess ΔAUC measure does not uniquely distinguish redundant from complementary information.

#### Synaptic-code visualization

To visualize the organization of task-related synaptic activity, small groups of spines were represented as binary activity configurations within the same 1-s epochs used for decoding. Each spine was assigned a binary value indicating the presence or absence of detected calcium activity during the corresponding epoch. Each resulting configuration represented the joint recruitment state of the selected spine group.

The frequencies of individual activity configurations were compared between epochs in which cursor error was improving and those in which error was worsening. These representations illustrated the distribution of joint synaptic recruitment patterns associated with task performance. Binary configurations were used for visualization and were distinct from the continuous event-rate predictors used to train the linear decoders and quantify functional synergy.

#### Relationship between functional synergy and plasticity

To investigate whether task-related synaptic interactions were associated with subsequent plasticity, excess synergy measured during early training (T1–T3) was related to changes in functional synaptic strength measured during late training (T6–T8). Associations were evaluated across spine groups of different sizes using Spearman rank correlations. To examine whether strongly synergistic and antagonistic groups differed in their subsequent plasticity, quadruplets were ranked according to their early-training excess synergy. The top and bottom 10% of the synergy distribution were designated positively synergistic and antagonistic groups, respectively. Within each group, plasticity polarity was quantified as the fraction of +Δw spines minus the fraction of -Δw spines.

Group-level plasticity polarity was compared between positively synergistic and antagonistic quadruplets. To account for the nonindependence introduced by overlapping spine groups, enrichment was evaluated using a spine-identity permutation procedure. Each spine was assigned a plasticity label (+1 for strengthened, −1 for weakened, and 0 for stable), and these labels were randomly permuted across eligible spines while preserving the overall distribution of plasticity classes. For each permutation, the shuffled labels were reassigned to the original quadruplets, preserving their membership and synergy rankings. The fractions of strengthened and weakened spines were recalculated for each group and averaged across positively synergistic and antagonistic groups separately. Plasticity enrichment was defined as the observed mean +Δw fraction minus the observed mean -Δw fraction, after subtracting the corresponding mean difference obtained from the shuffled distribution. This procedure was repeated 5,000 times for each analysis.

### Synaptic ensembles analyses

#### Event-centered synaptic population representations

For each eligible E1 dendritic event, simultaneously imaged spine activity was converted into a binary vector describing the participation of stable spine identities in the preceding recruitment interval. The principal ensemble definition was probabilistic: for each spine, its participation probability was estimated across relevant dendritic events, rather than enforcing a fixed top-k membership boundary. Participation was not restricted by interspine spatial distance.

#### Ensemble recurrence and recruitment selectivity

Event-to-event similarity of binary spine-recruitment vectors was used to quantify recurrence of synaptic patterns across learning. Recurrence was evaluated against an event-timing or circular-shift null preserving appropriate aspects of each spine’s activity statistics. The evolution of late-dominant spine identity, the fraction of recurrently recruited inputs, and enrichment or exclusion of strengthened and weakened identities were evaluated across training stages. In the main ensemble analysis, an event-centered pre-dendrite window of 1 s was used, with a two-frame ambiguity zone excluded near event onset.

#### Temporal focusing of synaptic recruitment

To assess changes in synaptic recruitment timing, dendrite-subtracted spine calcium-event activity was aligned to the onset of E1 dendritic calcium events. Analyses were restricted to events with a complete 4-s preceding recording window and no other E1 dendritic event within that interval. The final two imaging frames preceding dendritic-event onset were excluded to minimize ambiguity in event timing. The primary temporal-focusing analysis included all eligible E1 dendritic events, regardless of whether they were associated with successful E1 target acquisition. For analyses specifically examining reinforcement-associated recruitment, successful events were identified by matching E1 dendritic events to recorded E1 target acquisitions.

Temporal focusing was quantified as the normalized contrast between mean spine-event onset rates during the terminal (−1 to 0 s) and earlier (−3 to −1 s) intervals preceding dendritic-event onset:

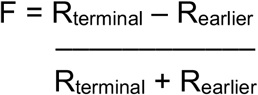

For each session, mean onset rates were first averaged across eligible dendritic events, and the normalized contrast was subsequently calculated from the pooled means. Positive values indicated greater recruitment during the final second preceding dendritic activation. The robustness of this measure was assessed using alternative earlier intervals of −4 to −1 s and −2 to −1 s.

#### Structural spine formation

Dendritic spine formation, elimination, and survival were quantified from longitudinal two-photon structural images using methods previously described^32^. The same dendritic segments were identified across imaging sessions using their morphology, spatial relationships to neighboring dendritic structures, and the positions of individual spines relative to stable anatomical landmarks. Individual dendritic protrusions extending at least one-third of the parent dendrite’s diameter were identified and tracked across the two baseline sessions (B2 and B1) and eight training sessions (T1–T8). Spine formation and elimination were determined by comparing corresponding dendritic segments across consecutive imaging sessions. Newly formed spines were defined as protrusions absent in the preceding session but present in the subsequent session, whereas eliminated spines were defined as protrusions present in the preceding session but absent in the subsequent session. Only dendritic segments with sufficient image quality to reliably resolve individual spines across imaging sessions were included. Imaging regions affected by substantial tissue displacement, rotation, motion artifacts, or insufficient structural visibility were excluded when these prevented unambiguous spine identification. Spines containing saturated fluorescence pixels were excluded, consistent with our previously established criteria^32^.

To characterize longitudinal structural fate, spines were assigned stable global identities and classified according to their presence across baseline and training. Stable spines were present during both baseline sessions and persisted through T8. Newly formed stable spines were absent during both baseline sessions, appeared during training, and remained present through T8. Newly formed transient spines were absent during baseline, appeared during training, but were subsequently eliminated before T8. Eliminated spines were present during both baseline sessions and subsequently disappeared, remaining absent throughout the remainder of training. Spines with intermittent, noncontiguous, or uncertain detection histories were classified as ambiguous and excluded from analyses requiring unambiguous structural fate.

Structural analyses were restricted to dendritic spines associated with the reinforced E1 dendrite. Newly formed spine recruitment into learned synaptic ensembles was evaluated by linking longitudinal structural identities to their corresponding functional calcium recordings and quantifying their activity and coactivation with pre-existing ensemble members during successful E1 events. Analyses specifically examining stabilized spine formation were restricted to newly formed spines that persisted through T8, whereas analyses of spine formation and subsequent survival distinguished stable from transient newly formed spines.

### Statistics and reproducibility

All analyses were performed using custom functions written in MATLAB 2024b (MathWorks). The independent biological unit was the animal, whereas dendrites, individual spines, dendritic calcium events, and sampled groups of spines constituted nested or repeated observations, as appropriate. These levels of observation were distinguished throughout the analyses to avoid treating technical subsampling as independent biological replication.

Longitudinal changes in behavioral performance, synaptic plasticity, and ensemble organization were assessed using repeated-measures analyses of variance (ANOVA) or linear mixed-effects models, as appropriate for the experimental design. For comparisons involving experimental treatment groups, models included treatment condition, training session or stage, and their interaction, with repeated measurements accounted for at the animal level. Planned comparisons between individual sessions, stages, or experimental conditions were performed using the corresponding paired or unpaired tests, as specified in the figure legends. Relationships between continuous variables were evaluated using Spearman rank correlations, ordinary least-squares regression, or multiple linear regression, depending on the hypothesis being tested.

For analyses involving shuffled-null distributions, statistical significance was assessed using permutation-based procedures tailored to the data structure. These included temporal shuffling of spine activity for coactivity and ensemble analyses, permutation of behavioral labels for decoding analyses, and permutation of spine identities or plasticity labels for analyses relating synaptic interactions to subsequent plasticity. Where applicable, permutations preserved the relevant experimental structure, including session membership, spine identity, or overlapping group membership. The number of permutations varied by analysis, typically ranging from 1,000 to 5,000, and is specified in the corresponding Methods subsections. Empirical permutation *p* values were calculated from the proportion of surrogate statistics at least as extreme as the observed statistic, using a finite-sample correction of adding one to both the numerator and denominator.

Multiple comparisons were corrected where appropriate. Unless otherwise indicated, statistical tests were two-sided, with a significance threshold of p < 0.05. Data are presented as mean ± s.e.m. unless otherwise specified; confidence intervals, medians, interquartile ranges, or individual observations are shown where appropriate. Exact sample sizes, definitions of independent experimental units, statistical tests, and p values are provided in the corresponding figure legends. * p < 0.05, ** p < 0.01, *** p < 0.001, ns: nonsignificant. No statistical methods were used to pre-determine sample sizes.

Animals, imaging sessions, dendritic segments, and spines were excluded according to predefined imaging-quality and data-completeness criteria described above. Imaging regions affected by substantial motion, insufficient structural visibility, or unreliable longitudinal spine identification were excluded.

#### Data and code availability

The source datasets supporting the figures, analysis scripts, and associated figure-generation code are available upon request and will be made publicly available upon publication in a peer-reviewed journal.

## Notes

### Competing Interest Statement

The authors have declared no competing interest.

## REFERENCES

1. Richards, B. A., & Lillicrap, T. P. (2019). Dendritic solutions to the credit assignment problem. Current opinion in neurobiology, 54, 28–36.

2. Körding, K. P., & König, P. (2001). Supervised and unsupervised learning with two sites of synaptic integration. Journal of computational neuroscience, 11(3), 207–215.

3. Magee, J. C., & Grienberger, C. (2020). Synaptic plasticity forms and functions. Annual review of neuroscience, 43(1), 95–117.

4. Lowet, A. S., Zheng, Q., Matias, S., Drugowitsch, J., & Uchida, N. (2020). Distributional reinforcement learning in the brain. Trends in neurosciences, 43(12), 980–997.

5. Roth, R. H., & Ding, J. B. (2024). Cortico-basal ganglia plasticity in motor learning. Neuron, 112(15), 2486–2502.

6. Economo, M. N., Komiyama, T., Kubota, Y., & Schiller, J. (2024). Learning and control in motor cortex across cell types and scales. The Journal of Neuroscience, 44(40), e1233242024.

7. Arber, S., & Costa, R. M. (2022). Networking brainstem and basal ganglia circuits for movement. Nature Reviews Neuroscience, 23(6), 342–360.

8. Peters, A. J., Liu, H., & Komiyama, T. (2017). Learning in the rodent motor cortex. Annual review of neuroscience, 40, 77–97.

9. Green, A. M., & Kalaska, J. F. (2011). Learning to move machines with the mind. Trends in neurosciences, 34(2), 61–75.

10. Sadtler, P. T., Quick, K. M., Golub, M. D., Chase, S. M., Ryu, S. I., Tyler-Kabara, E. C., … & Batista, A. P. (2014). Neural constraints on learning. Nature, 512(7515), 423–426.

11. Athalye, V. R., Santos, F. J., Carmena, J. M., & Costa, R. M. (2018). Evidence for a neural law of effect. Science, 359(6379), 1024–1029.

12. Clancy, K. B., Koralek, A. C., Costa, R. M., Feldman, D. E., & Carmena, J. M. (2014). Volitional modulation of optically recorded calcium signals during neuroprosthetic learning. Nature neuroscience, 17(6), 807–809.

13. Losey, D. M., Hennig, J. A., Oby, E. R., Golub, M. D., Sadtler, P. T., Quick, K. M., … & Chase, S. M. (2024). Learning leaves a memory trace in motor cortex. Current Biology, 34(7), 1519–1531.

14. Koralek, A. C., Jin, X., Long II, J. D., Costa, R. M., & Carmena, J. M. (2012). Corticostriatal plasticity is necessary for learning intentional neuroprosthetic skills. Nature, 483(7389), 331–335.

15. Francioni, V., Tang, V. D., Toloza, E. H., Brown, N. J., & Harnett, M. T. (2025). Vectorized instructive signals in cortical dendrites during a brain-computer interface task. bioRxiv, 2023–11.

16. Daie, K., Aitken, K., Rózsa, M., Bull, M. S., Humphreys, P. C., Wang, Z. C., … & Svoboda, K. (2026). Functional reorganization of motor cortex connectivity during learning. bioRxiv.

17. Cichon, J., & Gan, W. B. (2015). Branch-specific dendritic Ca2+ spikes cause persistent synaptic plasticity. Nature, 520(7546), 180–185.

18. Gonzalez, K. C., Negrean, A., Liao, Z., Terada, S., Zhang, G., Lee, S., … & Losonczy, A. (2025). Synaptic basis of feature selectivity in hippocampal neurons. Nature, 637(8048), 1152–1160.

19. Chen, Y., Kloos, M., Varga, Z., Zhang, Y., Piro, I., Sato, T. K., … & Konnerth, A. (2026). Thalamic activation of the visual cortex at the single-synapse level. Science, 391(6792), 1349–1354.

20. Yuste, R. (2011). Dendritic spines and distributed circuits. Neuron, 71(5), 772–781.

21. Frank, A. C., Huang, S., Zhou, M., Gdalyahu, A., Kastellakis, G., Silva, T. K., … & Silva, A. J. (2018). Hotspots of dendritic spine turnover facilitate clustered spine addition and learning and memory. Nature communications, 9(1), 422.

22. Kasai, H. (2023). Unraveling the mysteries of dendritic spine dynamics: Five key principles shaping memory and cognition. *Proceedings of the Japan Academy*, Series B, 99(8), 254–305.

23. Stuart, G. J., & Spruston, N. (2015). Dendritic integration: 60 years of progress. Nature neuroscience, 18(12), 1713–1721.

24. Kastellakis, G., & Poirazi, P. (2019). Synaptic clustering and memory formation. Frontiers in molecular neuroscience, 12, 300.

25. Scholl, B., & Fitzpatrick, D. (2020). Cortical synaptic architecture supports flexible sensory computations. Current opinion in neurobiology, 64, 41–45.

26. Kerlin, A., Mohar, B., Flickinger, D., MacLennan, B. J., Dean, M. B., Davis, C., … & Svoboda, K. (2019). Functional clustering of dendritic activity during decision-making. elife, 8, e46966.

27. Sehgal, M., Filho, D. A., Kastellakis, G., Kim, S., Lee, J., Shen, Y., … & Silva, A. J. (2025). Compartmentalized dendritic plasticity in the mouse retrosplenial cortex links contextual memories formed close in time. Nature Neuroscience, 28(3), 602–615.

28. Choi, D. I., Kim, J., Lee, H., Kim, J. I., Sung, Y., Choi, J. E., … & Kaang, B. K. (2021). Synaptic correlates of associative fear memory in the lateral amygdala. Neuron, 109(17), 2717–2726.

29. Kastellakis, G., Tasciotti, S., Pandi, I., & Poirazi, P. (2023). The dendritic engram. Frontiers in Behavioral Neuroscience, 17, 1212139.

30. Chidambaram, S. B., Rathipriya, A. G., Bolla, S. R., Bhat, A., Ray, B., Mahalakshmi, A. M., … & Sakharkar, M. K. (2019). Dendritic spines: Revisiting the physiological role. Progress in Neuro-Psychopharmacology and Biological Psychiatry, 92, 161–193.

31. Ma, S., & Zuo, Y. (2022, May). Synaptic modifications in learning and memory–A dendritic spine story. In Seminars in cell & developmental biology (Vol. 125, pp. 84–90). Academic Press.

32. Albarran, E., Raissi, A., Jáidar, O., Shatz, C. J., & Ding, J. B. (2021). Enhancing motor learning by increasing the stability of newly formed dendritic spines in the motor cortex. Neuron, 109(20), 3298–3311.

33. Stein, I. S., & Zito, K. (2019). Dendritic spine elimination: molecular mechanisms and implications. The Neuroscientist, 25(1), 27–47.

34. Hwang, F. J., Roth, R. H., Wu, Y. W., Sun, Y., Kwon, D. K., Liu, Y., & Ding, J. B. (2022). Motor learning selectively strengthens cortical and striatal synapses of motor engram neurons. Neuron, 110(17), 2790–2801.

35. Fu, M., Yu, X., Lu, J., & Zuo, Y. (2012). Repetitive motor learning induces coordinated formation of clustered dendritic spines in vivo. Nature, 483(7387), 92–95.

36. Egashira, R., Ke, M. T., Nakagawa-Tamagawa, N., Fujimoto, S., Inagaki, S., Takagi, T., … & Imai, T. (2026). Dendritic compartment-specific spine formation in layer 5 neurons underlies cortical circuit maturation during adolescence. Science Advances, 12(3), eadw8458.

37. Pandi, I., Chavlis, S., Oraby, H., Nashaat, M. A., Larkum, M., Papoutsi, A., & Poirazi, P. (2026). Dendritic hotspots support switching between competing rules without expanding the cortical engram. bioRxiv, 2026–08.

38. Hedrick, N. G., Lu, Z., Bushong, E., Singhi, S., Nguyen, P., Magaña, Y., … & Komiyama, T. (2022). Learning binds new inputs into functional synaptic clusters via spinogenesis. Nature neuroscience, 25(6), 726–737.

39. Harris, K. M. (2020). Synaptic odyssey. Journal of Neuroscience, 40(1), 61–80.

40. Dayan, P. (2009). Dopamine, reinforcement learning, and addiction. Pharmacopsychiatry, 42(S 01), S56–S65.

41. Gershman, S. J., & Uchida, N. (2019). Believing in dopamine. Nature Reviews Neuroscience, 20(11), 703–714.

42. Cox, J., & Witten, I. B. (2019). Striatal circuits for reward learning and decision-making. Nature Reviews Neuroscience, 20(8), 482–494.

43. Tang, J. C., Paixao, V., Carvalho, F., Silva, A., Klaus, A., da Silva, J. A., & Costa, R. M. (2024). Dynamic behaviour restructuring mediates dopamine-dependent credit assignment. Nature, 626(7999), 583–592.

44. Mohebi, A., Wei, W., Pelattini, L., Kim, K., & Berke, J. D. (2024). Dopamine transients follow a striatal gradient of reward time horizons. Nature Neuroscience, 27(4), 737–746.

45. Campbell, M. G., Ra, Y., Chen, Z., Xu, S., Burrell, M., Matias, S., … & Uchida, N. (2025). A hardwired neural circuit for temporal difference learning. bioRxiv.

46. Frémaux, N., & Gerstner, W. (2016). Neuromodulated spike-timing-dependent plasticity, and theory of three-factor learning rules. Frontiers in neural circuits, 9, 85.

47. Gerstner, W., Lehmann, M., Liakoni, V., Corneil, D., & Brea, J. (2018). Eligibility traces and plasticity on behavioral time scales: experimental support of neohebbian three-factor learning rules. Frontiers in neural circuits, 12, 53.

48. Wright, W. J., Hedrick, N. G., & Komiyama, T. (2025). Distinct synaptic plasticity rules operate across dendritic compartments in vivo during learning. Science, 388(6744), 322–328.

49. Yagishita S, Hayashi-Takagi A, Ellis-Davies GCR, Urakubo H, Ishii S, Kasai H. A critical time window for dopamine actions on the structural plasticity of dendritic spines. Science. 2014;345:1616–1620.

50. Fusi, S., Drew, P. J., & Abbott, L. F. (2005). Cascade models of synaptically stored memories. Neuron, 45(4), 599–611.

51. Richards, B. A., Lillicrap, T. P., Beaudoin, P., Bengio, Y., Bogacz, R., Christensen, A., … & Kording, K. P. (2019). A deep learning framework for neuroscience. Nature neuroscience, 22(11), 1761–1770.

52. Makino, H., Ren, C., Liu, H., Kim, A. N., Kondapaneni, N., Liu, X., … & Komiyama, T. (2017). Transformation of cortex-wide emergent properties during motor learning. Neuron, 94(4), 880–890.

53. Gilad, A., & Helmchen, F. (2020). Spatiotemporal refinement of signal flow through association cortex during learning. Nature communications, 11(1), 1744.

54. Yaeger, C. E., Vardalaki, D., Zhang, Q., Pham, T. L., Brown, N. J., Ji, N., & Harnett, M. T. (2024). A dendritic mechanism for balancing synaptic flexibility and stability. Cell reports, 43(8).

55. Neely, R. M., Koralek, A. C., Athalye, V. R., Costa, R. M., & Carmena, J. M. (2018). Volitional modulation of primary visual cortex activity requires the basal ganglia. Neuron, 97(6), 1356–1368.

56. Vendrell-Llopis, N., Read, J., Boggiano, S., Hetzler, B., Peitsinis, Z., Stanley, C., … & Isacoff, E. Y. (2025). Dopamine D1 receptor activation in the striatum is sufficient to drive reinforcement of anteceding cortical patterns. Neuron, 113(5), 785–794.

57. Hosp, J. A., & Luft, A. R. (2013). Dopaminergic meso-cortical projections to M1: role in motor learning and motor cortex plasticity. Frontiers in neurology, 4, 145.

58. Ghanayim, A., Benisty, H., Cohen Rimon, A., Schwartz, S., Dabdoob, S., Lifshitz, S., … & Schiller, J. (2025). VTA projections to M1 are essential for reorganization of layer 2-3 network dynamics underlying motor learning. Nature Communications, 16(1), 200.

59. Audette, N. J., & Schneider, D. M. (2023). Stimulus-specific prediction error neurons in mouse auditory cortex. The Journal of Neuroscience, 43(43), 7119–7129.

60. Jain, A., Nakahata, Y., Pancani, T., Watabe, T., Rusina, P., South, K., … & Yasuda, R. (2024). Dendritic, delayed, stochastic CaMKII activation in behavioural time scale plasticity. Nature, 635(8037), 151–159.

61. Chen, S. X., Kim, A. N., Peters, A. J., & Komiyama, T. (2015). Subtype-specific plasticity of inhibitory circuits in motor cortex during motor learning. Nature neuroscience, 18(8), 1109–1115.

62. Pi, H. J., Hangya, B., Kvitsiani, D., Sanders, J. I., Huang, Z. J., & Kepecs, A. (2013). Cortical interneurons that specialize in disinhibitory control. Nature, 503(7477), 521–524.

63. Szadai, Z., Pi, H. J., Chevy, Q., Ócsai, K., Albeanu, D. F., Chiovini, B., … & Rózsa, B. (2022). Cortex-wide response mode of VIP-expressing inhibitory neurons by reward and punishment. Elife, 11, e78815.

64. He K, Huertas M, Hong SZ, Tie X, Hell JW, Shouval H, Kirkwood A. Distinct eligibility traces for LTP and LTD in cortical synapses. Neuron. 2015;88:528–538.

65. Hangya, B., Ranade, S. P., Lorenc, M., & Kepecs, A. (2015). Central cholinergic neurons are rapidly recruited by reinforcement feedback. Cell, 162(5), 1155–1168.

66. Grossman, C. D., & Cohen, J. Y. (2022). Neuromodulation and neurophysiology on the timescale of learning and decision-making. Annual review of neuroscience, 45, 317–337.

67. Yu, B., Yue, Y., Ren, C., Yun, R., Lim, B., & Komiyama, T. (2025). Cholinergic feedback for modality-and context-specific modulation of sensory representations. Science, 388(6753), 1324–1329.

68. Francioni, V., & Harnett, M. T. (2022). Rethinking single neuron electrical compartmentalization: dendritic contributions to network computation in vivo. Neuroscience, 489, 185–199.

69. Landau, A. T., Park, P., Wong-Campos, J. D., Tian, H., Cohen, A. E., & Sabatini, B. L. (2022). Dendritic branch structure compartmentalizes voltage-dependent calcium influx in cortical layer 2/3 pyramidal cells. Elife, 11, e76993.

70. Wong-Campos, J. D., Park, P., Lee, B. H., Davis, H. C., Qi, Y., Tian, H., … & Cohen, A. E. (2026). Voltage dynamics of cortical dendrites in vivo. Nature Neuroscience, 1–9.

71. Otor, Y., Achvat, S., Cermak, N., Benisty, H., Abboud, M., Barak, O., … & Schiller, J. (2022). Dynamic compartmental computations in tuft dendrites of layer 5 neurons during motor behavior. Science, 376(6590), 267–275.

72. O’Hare, J. K., Gonzalez, K. C., Herrlinger, S. A., Hirabayashi, Y., Hewitt, V. L., Blockus, H., … & Losonczy, A. (2022). Compartment-specific tuning of dendritic feature selectivity by intracellular Ca2+ release. Science, 375(6586), eabm1670.

73. Zhang, Y., Liu, Y., Hu, Y., Zhang, Z., Du, J., Cui, L., … & Xu, N. L. (2026). Dendritic computation for abstract rule-based flexible decision-making. Science, 394(6821), eadz5323.

74. Noguchi, A., Terada, S., Zakka, G. N., O’Toole, C., Reynolds, L., Land, M. A., … & Losonczy, A. (2026). Parallel independent voltage computing along dendrites of CA3 pyramidal neurons. Science, 393(6808), 306–312.

